# A genetic selection reveals functional metastable structures embedded in a toxin-encoding mRNA

**DOI:** 10.1101/615682

**Authors:** Sara Masachis, Nicolas J. Tourasse, Claire Lays, Marion Faucher, Sandrine Chabas, Isabelle Iost, Fabien Darfeuille

**Author notes:** Faculty of Biology I, Department of Microbiology, Ludwig-Maximilians-University of Munich, 82152 Martinsried, Germany (Sara Masachis).

## Abstract

Post-transcriptional regulation plays important roles to finely tune gene expression in bacteria. In particular, regulation of type I toxin-antitoxin (TA) systems is achieved through sophisticated mechanisms involving toxin mRNA folding. Here, we set up a genetic approach to decipher the molecular underpinnings behind the regulation of a type I TA in *Helicobacter pylori*. We used the lethality induced by chromosomal inactivation of the antitoxin to select mutations that suppress toxicity. We found that single point mutations are sufficient to allow cell survival. Mutations located either in the 5’ untranslated region or within the open reading frame of the toxin hamper its translation by stabilizing stem-loop structures that sequester the Shine-Dalgarno sequence. We propose that these short hairpins correspond to metastable structures that are transiently formed during transcription to avoid premature toxin expression. This work uncovers the co-transcriptional inhibition of translation as an additional layer of TA regulation in bacteria.

## INTRODUCTION

In any living cell, unwanted gene expression can have a detrimental effect on cell growth, and eventually lead to cell death. In bacteria, a fine tuning of gene expression can be achieved at the translational level through the control of the ribosome binding site (RBS) accessibility, which encompasses the Shine-Dalgarno (SD) sequence and the start codon. Messenger RNA structures occluding the RBS have been reported to control the expression of many important genes for which a timely control is crucial. Regulation of translation initiation via SD-sequestration is an old theme that initially started with the study of bacteriophage genes (De Smit & Duin, 1990) and ribosomal proteins (for review see Duval et al., 2015). More recently, its impact on other bacterial genes such as sigma factors (Mearls et al., 2018) and translational riboswitches (Rinaldi, Lund, Blanco, & Walter, 2016) has been reported.

Hence, in many cases, preventing gene expression via SD-sequence sequestration is crucial. This is particularly true for type I toxin-antitoxin (TA) systems. In contrast to the largest type II TA family, antitoxins belonging to type I TA systems do not directly interact with the toxin protein but rather prevent its expression (Harms, Brodersen, Mitarai, & Gerdes, 2018). This regulation occurs through the direct base-pairing of the RNA antitoxin with the toxin mRNA and leads to toxin translation inhibition and/or mRNA degradation (Brantl & Jahn, 2015; Durand, Jahn, Condon, & Brantl, 2012; Wen & Fozo, 2014). However, the action of the RNA antitoxin is often not sufficient to avoid toxin expression (Masachis & Darfeuille, 2018). Indeed, due to the coupling between transcription and translation occurring in bacteria, translation of the nascent toxin mRNA can potentially occur before antitoxin action. Thus, a tight control of toxin synthesis is usually achieved via the direct sequestration of its SD sequence within stable stem-loop structures (Masachis & Darfeuille, 2018). The existence of a non-translatable toxin primary transcript is a major hallmark of type I TA loci. In this transcript, the RBS occlusion occurs through the base-pairing of the SD sequence with a partially or totally complementary sequence termed anti-SD (aSD). Such aSD sequences are often located a few nucleotides (up to 11) upstream or downstream of the SD sequence and trap it within a hairpin structure. However, in some cases, RBS occlusion involves an aSD sequence encoded far downstream and is achieved via a long-distance interaction (LDI) between the 5’ and 3’ ends of the toxin mRNA (Gultyaev, Franch, & Gerdes, 1997; Han, Kim, Bak, Park, & Lee, 2010).

This strategy of toxin expression regulation via the formation of a LDI has been recently described for a type I TA family of the *Epsilon* proteobacteria. This family, named *aapA*/IsoA, is present in several copies on the chromosome of the major human gastric pathogen *Helicobacter pylori*. We characterized the *aapA1*/IsoA1 TA system at the locus I and showed that the *aapA1* gene codes for a small toxic protein whose expression is repressed by a *cis*-overlapping antisense RNA, IsoA1 (Arnion et al., 2017). We have shown that transcription of this toxic gene generates a highly stable primary transcript whose translation is post-transcriptionally impeded by a 5’-3’ LDI. Consequently, a 3’-end ribonucleolytic event, that we termed ‘mRNA activation step’, is necessary to remove the LDI, thus enabling toxin translation (Arnion et al., 2017).

In the present work, we aimed at deciphering the mechanism of expression of another module belonging to the *aapA*/IsoA family, the *aapA3*/IsoA3. We first showed that, in the absence of antitoxin expression, chromosomal toxin expression is lethal. Taking advantage of this lethal phenotype, we used a genetic approach to select suppressors allowing survival. This method, that we previously named FASTBAC-Seq for Functional AnalysiS of Toxin-antitoxin in BACteria by deep Sequencing, allows the mapping of intragenic toxicity suppressors within a given TA locus (Masachis, Tourasse, Chabas, Bouchez, & Darfeuille, 2018). In the case of the *aapA3*/IsoA3 TA locus, FASTBAC-Seq revealed that single point mutations are sufficient to counteract the lethality caused by the absence of antitoxin. Unexpectedly, one-third of suppressors mapped to non-coding regions of the toxin mRNA. Some of them target well-known regulatory elements such as the toxin promoter and the SD sequence. Remarkably, we showed that one of the suppressors located in the SD sequence does not act at the sequence level but at the mRNA structural level. Indeed, this mutation inhibits translation of the active mRNA by stabilizing an RNA hairpin in which the SD sequence is masked by an upstream-encoded aSD sequence (aSD1). A synonymous mutation within the Open Reading Frame (ORF) acts similarly but on another hairpin in which the SD sequence is masked by a downstream-encoded aSD (aSD2). These suppressor mutations reveal two transient hairpin structures that sequentially form during transcription and which are then replaced by a more stable LDI upon transcription termination. Our results indicate that, in addition to the post-transcriptional control achieved via a stable LDI, metastable structures are also required to prevent premature toxin expression in a co-transcriptional manner.

## RESULTS

### The small antisense RNA IsoA3 is essential to prevent AapA3 translation

We previously studied the regulation of *aapA1*/IsoA1, a member of the *aapA*/IsoA type I TA family recently identified in *H. pylori* (Arnion et al., 2017). Here, we studied the *aapA3*/IsoA3 module (Figure 1A; for sequence details see Figure 1 – figure supplement 1). As other TA systems of this family, the *aapA3*/IsoA3 locus codes for an antisense RNA, IsoA3 (80 nucleotides), encoded on the opposite strand of a small ORF, AapA3. The AapA3 peptide (30 amino acids) shares 60% sequence homology with the AapA1 peptide, whose ectopic expression is toxic in *H. pylori* (Arnion et al., 2017). Here, we first investigated whether *aapA3* expression from the chromosome is toxic. For this purpose, we inactivated the antitoxin promoter by introducing two point mutations in its −10 box, while maintaining the amino acid sequence of the toxin (Figure 1B, pIsoA3* in all figures), as previously described for the IsoA1 promoter (Arnion et al., 2017). Insertion of these mutations on the chromosome was performed using a counter-selection cassette, which allows the generation of unmarked transformants (Dailidiene, Dailide, Kersulyte, & Berg, 2006). Briefly, the TA locus of a streptomycin resistant 26695 strain (K43R) was replaced by the *rpsl_Cj_-erm* double marker cassette, giving rise to the streptomycin sensitive Δ*aapA3*/IsoA3*::rpsl_Cj_-erm*/K43R strain (see Figure 3B) (Masachis et al., 2018). Then, we performed gene replacement assays using PCR constructs carrying either a wild-type or an antitoxin inactivated (pIsoA3*) TA locus. Strains that had undergone homologous recombination were selected on streptomycin. However, no transformants were obtained unless a non-sense or a frameshift mutation was introduced in the *aapA3* ORF. This result indicated that, in the absence of IsoA3 synthesis, the AapA3 toxin expression from its chromosomal locus is constitutive and too toxic to obtain a viable strain.

**Figure 1.**
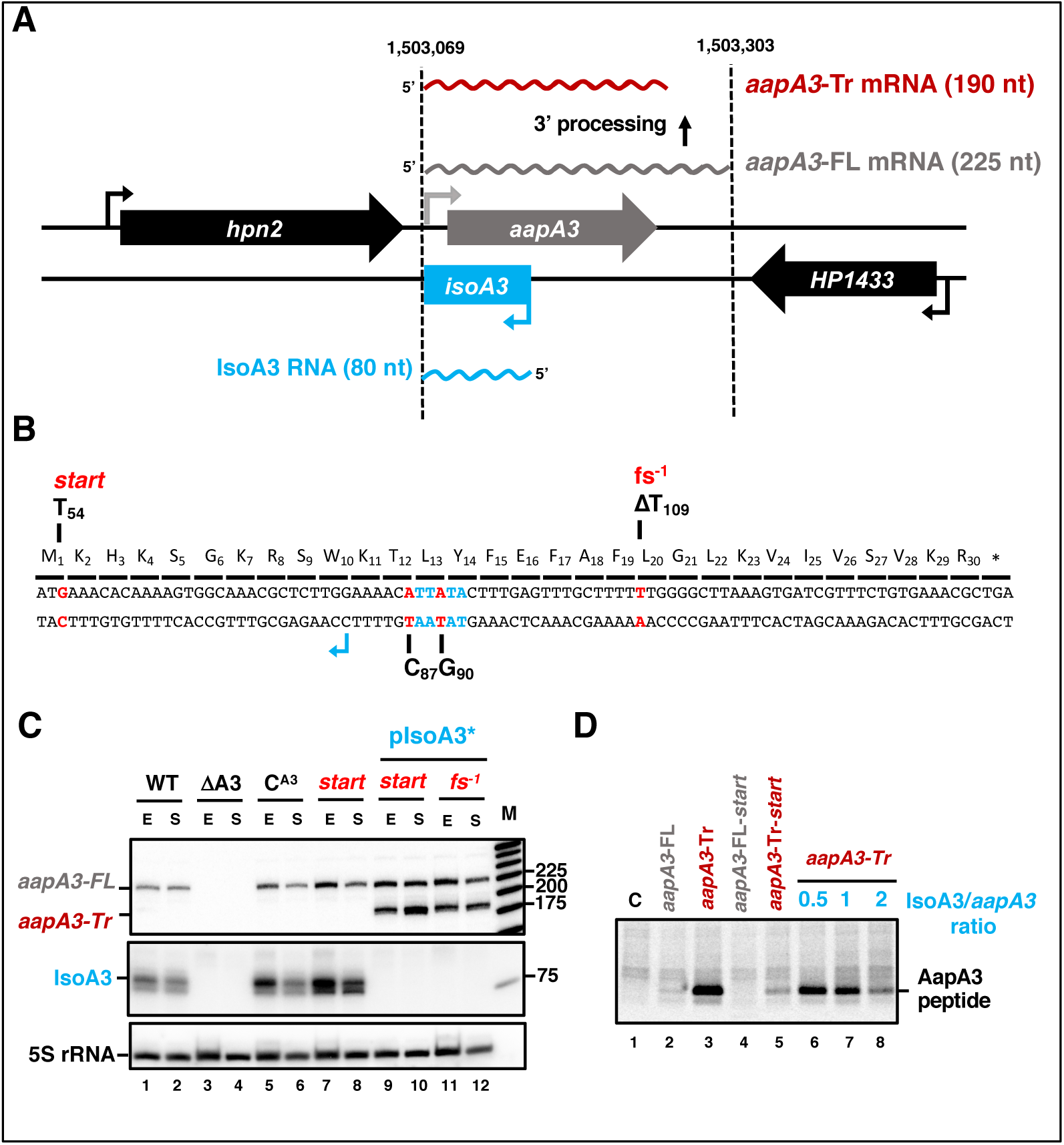
IsoA3 small RNA is essential to prevent AapA3 translation. (A) Organization of the aapA3/IsoA3 locus in the *H. pylori* 26695 strain. Grey arrow, AapA3 ORF; blue box, IsoA3 RNA; small bent arrows, −10 box of each transcript. Grey, red and blue wavy lines represent *aapA3*-FL (full-length), *aapA3*-Tr (3’-end truncated) and IsoA3 transcripts, respectively. Their approximate length is also indicated. (B) Nucleotide and amino acid sequence of AapA3 ORF with hallmarks. The sequence of the IsoA3 promoter (−10 box) is shown in blue. The nucleotides that were mutated to inactivate the IsoA3 promoter (pIsoA3*) and the AapA3 start codon (start), and to create a −1 frameshift (fs-1) are shown in red. (C) The ‘WT’ strain corresponds to the 26695 *H. pylori* strain containing the K43R mutation in the *rpsL* gene, which confers resistance to streptomycin. The ΔA3 strain is the parental strain in which the aapA3/IsoA3 locus has been replaced by the *rpsL*_Cj_-erm cassette (ΔaapA3/IsoA3::*rpsL*_Cj_-erm). The C^A3^ and start strains correspond to the ΔA3 strain complemented with the WT aapA3/IsoA3 locus and with the locus mutated at the start codon (G54T), respectively. The two strains inactivated for the IsoA3 promoter (pIsoA3*) also contain a frameshift mutation (fs-1) or a mutation in the start codon (start). Total RNA from stationary (S) or exponential (E) growth phase of the indicated strains was isolated and subjected to Northern blot. The aapA3-FL, aapA3-Tr, and IsoA3 transcripts are shown. 5S rRNA assessed proper loading. (D) Translation assays were performed with 0.5 µg of *aapA3* mRNAs in absence or presence of IsoA3, in 0.5, 1 or 2 molar ratios. [^35^S]-Met was used for labeling. The control lane (C) shows the translation background obtained without exogenous mRNA.

Total RNA of two viable transformants containing either a mutation at the start codon (*start*) or a −1 frameshift mutation leading to a premature stop codon (*fs^−1^*) (Figure 1B) were analyzed by Northern Blot (Figure 1C). The absence of IsoA3 transcript (Figure 1C, lanes 9-12) confirmed the successful inactivation of the IsoA3 promoter. As a control, the complementation of the Δ*aapA3*/IsoA3*::rpsl_Cj_-erm*/K43R strain (ΔA3) with the WT *aapA3*/IsoA3 locus (C^A3^) was successfully achieved with no change in the expression pattern (Figure 1C, compare lanes 5-6 with lanes 1-2). Two *AapA3* mRNA species were detected in absence of IsoA3 expression (Figure 1C, lanes 9 to 12): a long transcript of 225 nt, which was denoted *aapA3*-FL (full-length) and a shorter transcript lacking the last 35 nt (*aapA3*-Tr). This latter was not detected in presence of IsoA3 RNA (Figure 1C, lanes 1, 2, 5-8) and corresponds to the truncated mRNA species previously described for the *aapA1*/IsoA1 homolog (Arnion et al., 2017). *In vitro* translation assays demonstrated that only the truncated mRNA is efficiently translated (Figure 1D, lane 3). Translation of *aapA3*-FL (Figure 1D, lane 2) or of the *aapA3*-FL containing a non-sense mutation in the start codon (Figure 1D, lane 4) were close to the translational background obtained in absence of exogenous mRNA (Figure 1D, lane 1). Importantly, the absence of IsoA3 leads to the accumulation of *aapA3*-Tr without affecting the amount of *aapA3*-FL (Figure 1C lanes 9-12), indicating that IsoA3 specifically targets *aapA3*-Tr *in vivo*. *In vitro* structure probing of the two AapA3 mRNA species further confirmed that IsoA3 only interacts with the *aapA3*-Tr mRNA (Figure 2, lanes 4-7). Base-pairing between both transcripts creates an extended RNA heteroduplex of 80 base-pairs (Figure 2, lane 4) that is translationally inert, as shown by *in vitro* translation assays (Figure 1D, lanes 6-8). Remarkably, none of the IsoA RNAs produced from the five other *aapA*/IsoA chromosomal loci (I, II, IV, V and VI) can replace the absence of IsoA3 expression demonstrating that their regulation is strictly module-specific, as previously suggested by *in vitro* translation assays (Sharma et al. 2010).

**Figure 2.**
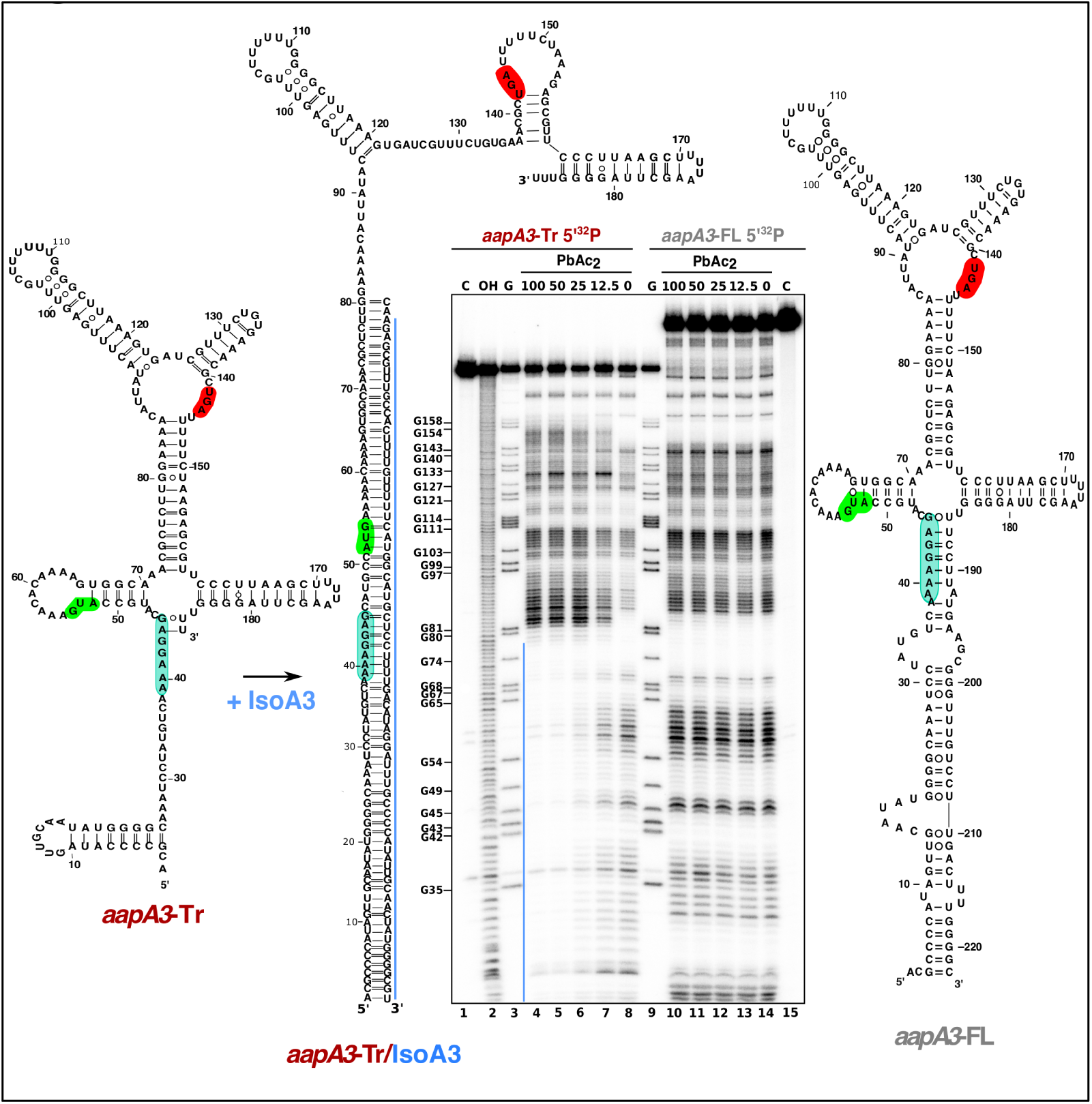
IsoA3 inhibits *aapA3*-Tr translation by masking its SD region. ∼0.1 pmol of 5’-end [^32^P]-labeled *in vitro* transcribed *aapA3-*FL and *aapA3*-Tr RNAs were subjected to lead probing in presence of increasing concentrations of IsoA3 (0-100 nM). Untreated RNA (lanes 1 and 15, denoted C) and partially alkali digested RNA (denoted OH, lane 2) served as control and ladder, respectively. Positions of all G residues revealed upon T1 digestion under denaturing conditions (lanes ‘G’) are indicated relative to the transcription start site of the *aapA3* gene. Cleaved fragments were analyzed on an 8% denaturing PAA gel. 2D structure predictions were generated with the RNAfold Web Server (Gruber, Lorenz, Bernhart, Neuböck, & Hofacker, 2008) and VARNA (Darty, Denise, & Ponty, 2009) was used to draw the diagrams. The region involved in duplex formation between IsoA3 and *aapA3*-Tr mRNA is indicated with a blue line; the start codon, stop codon and SD sequence of AapA3 are highlighted in green, red and turquoise, respectively.

Altogether, we showed that IsoA3 represses *aapA3* constitutive expression at the translational level by forming a stable RNA heteroduplex. This duplex is then likely degraded by the double-stranded specific ribonuclease RNase III, leading to a rapid turnover of the translationally active toxin-encoding mRNA, as shown for the *aapA1*/IsoA1 locus (Arnion et al. 2017).

### Decoding AapA3 toxicity determinants with nucleotide resolution

To identify the toxicity determinants of this TA locus, we next aimed at taking advantage of the lethality induced by the chromosomal inactivation of the IsoA3 antitoxin to search for toxicity suppressors. To this end, we performed the same gene replacement assays as described above, using two PCR constructs carrying either a wild-type TA locus (WT) or two synonymous mutations inactivating the IsoA3 promoter (pIsoA3*). Two PCR containing either a point mutation inactivating the start codon (*start*) or both mutations (*start*/pIsoA3*) were also used as controls (Figure 3A). These four PCR constructs were transformed into the Δ*aapA3*/IsoA3*::rpsl_Cj_-erm*/K43R *H. pylori* strain and Str^R^ transformants were selected on streptomycin-containing plates. For each transformation, the number of streptomycin-resistant colonies was determined and normalized to the total number of transformed cells (Figure 3B). As expected, when no DNA was used for transformation (H_2_O) only phenotypic revertants having mutated the *rpsl_Cj_* gene were selected (Masachis et al., 2018). The lack of IsoA3 expression led to a strong reduction (1.83 log-fold) in the number of transformed cells compared to that obtained with the WT or double-mutant *start*/pIsoA3* constructs (Figure 3B). These results confirmed that in the absence of IsoA3, the chromosomal expression of AapA3 is highly toxic. Remarkably, the number of transformants obtained in absence of antitoxin was slightly higher to that obtained when no DNA was used for transformation (Figure 3B). The sequencing of the TA locus of approximately hundred pIsoA3* transformants revealed that all of them contained point mutations in the AapA3 ORF (Masachis et al., 2018). Thus, our genetic approach selected mutations that suppress toxicity allowing the generation of recombinant strains lacking antitoxin expression.

**Figure 3.**
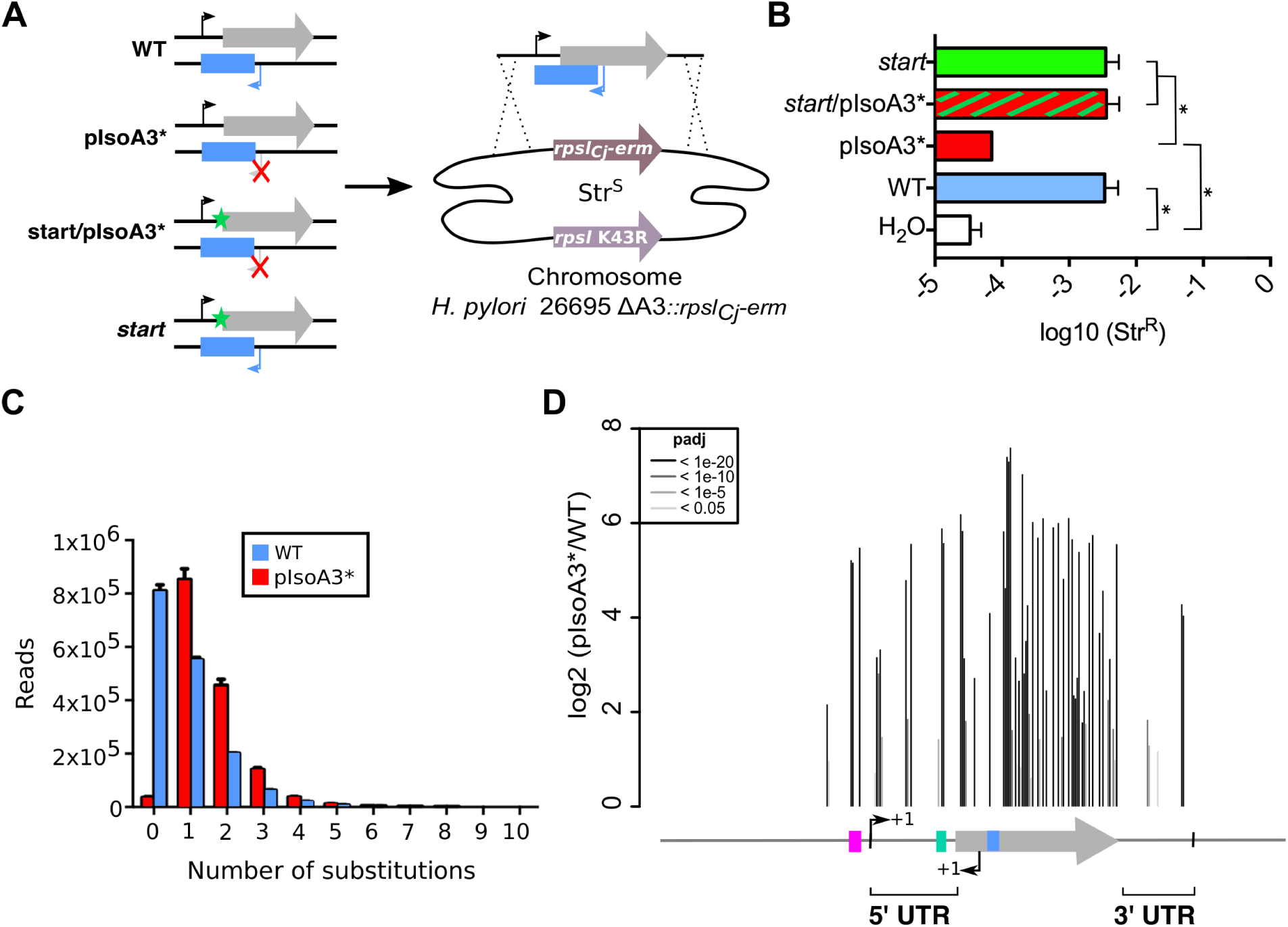
Unveiling intragenic toxicity determinants with nucleotide resolution. **(A)** PCR fragments used for transformation of the ΔA3 strain (Δ*aapA3*/IsoA3::*rpsL_Cj_-erm)* are shown. A green star indicates a mutation in the start codon (G54T). The red cross indicates the two mutations (T87C and T90G) introduced to inactivate the IsoA3 promoter (pIsoA3*). To select recombination at the locus, Str^R^ transformants were selected. **(B)** Transformation efficiency (number of Str^R^ transformants divided by the total number of transformed cells) was determined for each construct. A control in which the PCR fragment was replaced by H_2_O is also shown. Error bars represent standard deviations (s.d); *n=3* biological replicates (\**P*<0.05; values according to unpaired *t*-test). **(C)** Number of reads containing 0 to 10 substitutions in the sequenced amplicon of 426 nt encompassing the *aapA3*/IsoA3 TA locus. Error bars represent s.d; *n=3* biological replicates. **(D)** Positional analysis of single-nucleotide substitutions on the *aapA3*/IsoA3 locus. Bar plot shows the log2 fold change (pIsoA3*/WT ratio) for the 70 positions with an adjusted *p*-value (padj) lower than 0.05. Bars are drawn with different shades of grey according to the *p*-value. The relevant sequence elements are indicated by arrows and boxes under the graph. 5’ UTR, 5’ untranslated region; purple box, *aapA3* −10 box; small bent arrows, +1 transcription start site (TSS) of *aapA3* and IsoA3; turquoise box, *aapA3* SD sequence; large grey arrow, *aapA3* ORF; small blue box, IsoA3 −10 box; 3’ UTR, 3’ untranslated region.

To explore the complete landscape of suppressors, we next scaled-up the transformation assay using the WT or pIsoA3* PCR products as DNA substrates. The transformation assay was performed using three independent biological replicates for each construct. Approximately 60,000 transformants per construct were collected and pooled, genomic DNA was extracted, and an amplicon of 426 nt encompassing the whole TA locus was sequenced by the Illumina paired-end approach. This approach, called FASTBAC-Seq, has been described recently (Masachis et al., 2018). Consistent with the above-mentioned first transformation assay, deep-sequencing data analysis showed that 97.7% of the pIsoA3* transformants contained mutations (Figure 3C). A strong mutation rate (51.2%) was also observed with the WT PCR product (Figure 3C). This result was explained by an unanticipated technical artifact linked to the PCR assembly, which led to a strong mutation rate in the overlapping region used for assembly (nucleotides 80, 81 and 82 of *aapA3*, the +1 corresponding to the transcription start site [TSS]). This artifact strongly reduced the sequencing depth and impeded the analysis of double and triple mutations. Consequently, we focused our analysis on single-nucleotide mutations that have been significantly enriched (adjusted False Discovery Rate padj ≤ 0.05) in absence of antitoxin relative to WT (Figure 3D).

Analysis of the number of mutations per read in the complete sequencing dataset (all replicates combined) showed that more than half of the pIsoA3* transformed clones (51.8% out of ∼5.1 million) were mutated at a single nucleotide position (Figure 3C). This result demonstrates that single point mutations are sufficient to abolish AapA3 toxin activity and/or expression. A low number of pIsoA3* strains (2.3%) had a wild-type locus sequence (Figure 3C, pIsoA3* zero mutation), which can be explained by the sequencing error rate (around 1.5%) and/or suppressor mutations lying in regions outside the TA locus (*i.e.*, outside the amplicon). Single-nucleotide mutations were mainly substitutions, which are favored by PCR biases. Only 4% were insertions and deletions (indels). Hierarchical clustering analysis revealed that the location and frequency of single substitutions were highly similar in the three biological replicates, indicating that the locus coverage was close to optimum (Figure 3 – figure supplement 1). Contrary to substitutions (which were found in both coding and non-coding regions), the single-nucleotide indels were almost exclusively present in the AapA3 ORF, generating truncated or longer forms of the peptide. Moreover, in some cases, they were not present in all three replicates, reflecting their under-representation in the PCR products (Figure 3 – figure supplement 1). Importantly, depending on the type of statistical analysis, “position-specific” or “nucleotide-specific” (see Material and Methods for details), there were statistically enriched substitutions in absence of antitoxin (padj ≤ 0.05) at 70 or 72 positions within the *aapA3*/IsoA3 locus, respectively (65 positions in common between the two analyses). Substitutions identified in only one of the analyses included (relative to *aapA3* +1 TSS): i) positions −26 and −7 within the promoter region; ii) position +28 in the 5’ UTR; iii) positions +64 and +97 in the AapA3 ORF; and iv) positions +146 and +177 in the 3’ UTR. Such positions had generally a close-to-cut-off padj value, but not in all cases. For instance, position +28, which has been studied hereafter, had a highly significant padj value of 7.2×10^−6^.

Expectedly, the highest mutation densities (defined as the number of mutated nucleotides divided by the total number of nucleotides in the region of interest) were observed in the AapA3 ORF (53%), as well as in well-known regulatory regions such as the −10 box of the toxin mRNA (66.7%, Figure 3 – figure supplement 2) and the SD sequence (42.8%) (Figure 3D). For the *aapA3* −10 box, out of the six positions (5’-TAGGAT-3’), suppressor mutations were mostly found in the first two and last nucleotides (Figure 3 – figure supplement 2). This result allowed us to determine the minimal functional *aapA3* −10 box motif 5’-TANNNT-3’, which is in perfect agreement with the previously determined consensus sequence (C. M. Sharma et al., 2010). This result also confirmed that the arbitrarily-chosen False Discovery Rate cut-off (padj ≤ 0.05) was stringent enough to avoid false positives but permissive enough to allow the identification of suppressor substitutions. Remarkably, seventeen mutations were unveiled in the 5’ and 3’ untranslated regions (Figure 3D). In the present work, we have focused our study to mutations lying around the RBS.

### Two suppressor mutations in the 5’ UTR impede AapA3 translation

The genetic selection of suppressors allowed us to determine the minimal functional sequence of the toxin SD sequence. Among the seven positions (5’-AAAGGAG-3’), substitutions at the two central guanine nucleotides (positions +42 and +43) were the most frequently mutated in absence of antitoxin (Figure 4A). As expected from PCR biases (Beaudry & Joyce, 1992), although the transition G>A was preferentially enriched in both cases, the G>C and G>T transversion mutations were also selected. Strikingly, a less-frequent transversion mutation (A>T) was selected within the SD sequence at position +40 (A40T, Figure 4A). Moreover, another unique transversion upstream of the SD sequence was identified (A28C, Figure 4A). Strains containing these atypical mutations were constructed and further analyzed.

**Figure 4.**
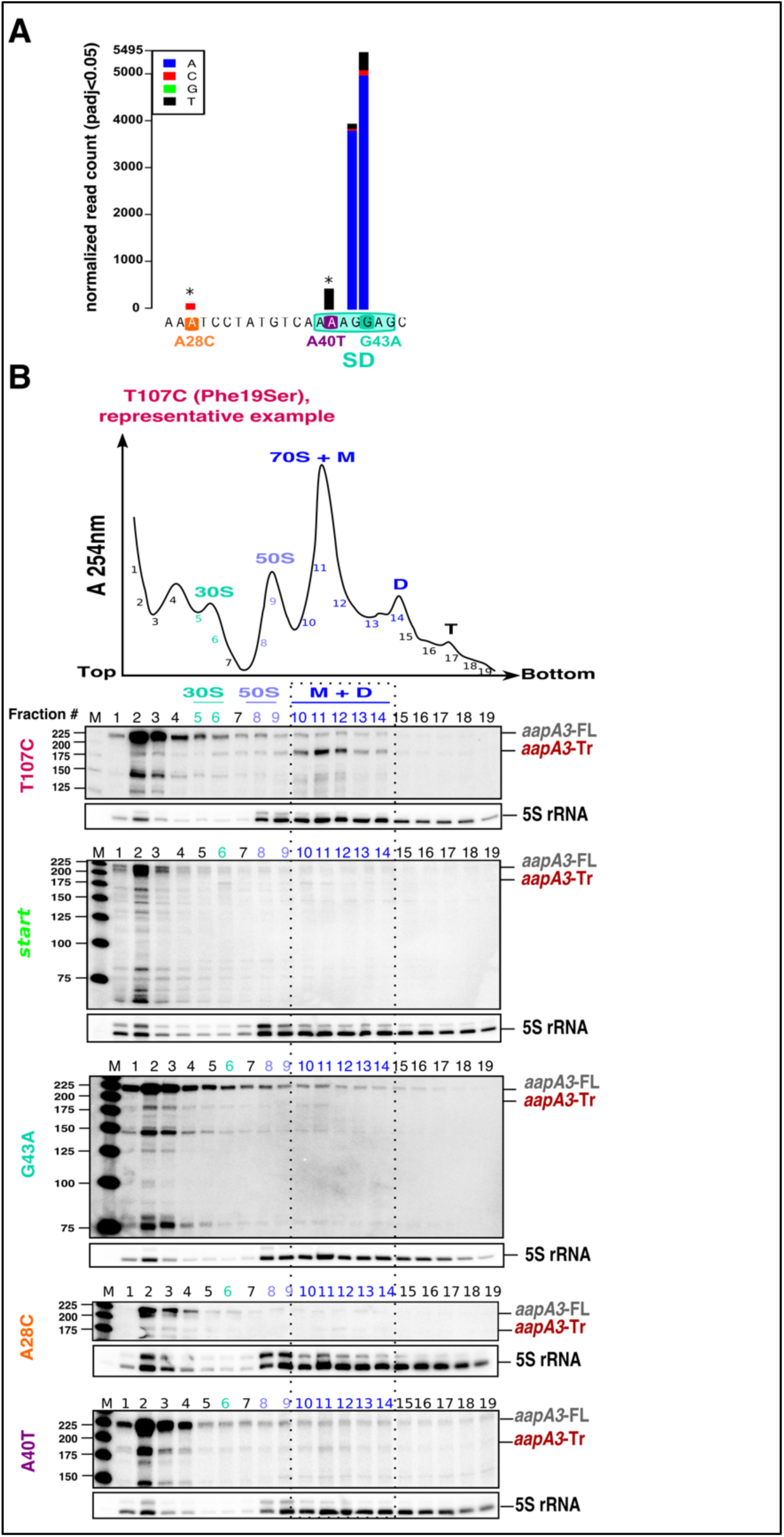
Single point suppressor mutations in the 5’ UTR of the *aapA3* mRNA inhibit its translation. **(A)** Nucleotide substitutions within the *aapA3* 5’ UTR, which are significantly enriched (pajd≤0.05, “nucleotide-specific” analysis, see Material and Methods section) in pIsoA3* compared to WT. Asterisks above the bars indicate transversion mutations. The SD sequence is boxed. **(B)** Cell lysates of the indicated *aapA3* variant strains were subjected to ultracentrifugation through a sucrose gradient. A representative A_254nm_ profile of the T107C strain is shown. Peaks of the free 30S and 50S subunits, 70S ribosomes (free ribosomes and monosomes (M)), and polysomes (D, disomes; T, trisomes) are indicated. RNA was extracted from each fraction and equal volumes of each fraction were subjected to Northern blot analysis. The different transcripts *aapA3*-FL, *aapA3*-Tr, and 5S rRNA (loading control) are indicated. The vertical dashed lines delineate the limits corresponding to 70S, monosome and disome fractions.

Due to their proximity to the SD sequence, we first tested whether these mutations could affect AapA3 translation *in vivo*. Due to the lack of AapA3-targeting antibodies, its translation was assessed indirectly by polysome fractionation coupled to Northern blot analysis. As a control, we used a suppressor mutation isolated during our FASTBAC-Seq selection, which inactivates the toxin activity but not its expression. We thus generated a strain containing the pIsoA3* mutations in combination with a mutation in the toxin ORF converting a phenylalanine at position 19 into a serine (T107C, Figure 4B). Polysome fractionation of this strain confirmed that the toxin full-length mRNA (*aapA3-FL*) is mainly found in non-ribosomal fractions (only 3% was present in polysomes, lanes 10 to 14 in Figure 4B; see Figure 4 – figure supplement 2 for quantification) whereas the truncated isoform (*aapA3*-Tr) is associated with the monosome and disome fractions (73% present in these fractions; Figure 4B and Figure 4 – figure supplement 2A). The absence of the *aapA3*-Tr form in heavier polysomes is probably due to the short length of the ORF (90 nucleotides), which cannot accommodate more than two ribosomes. This result clearly confirmed that the *aapA3*-FL is indeed a translationally inert isoform whereas the *aapA3*-Tr is translationally active. Hence, polysome fractionation is a powerful tool to study translation efficiency of toxin mRNAs *in vivo*.

We then analyzed the efficiency of toxin mRNA translation in strains containing a mutation either in the start codon (*start*) or in the SD sequence (G43A) (Figure 4B). In both cases, the active mRNA isoform (*aapA3*-Tr) was no more found within the polysome fractions. Instead, significant levels of *aapA3*-Tr mRNA degradation products were detected in the top of the gradient, which may arise from the lack of ribosome protection and/or the extended time of sample collection and treatment prior RNA extraction. The strong degradation observed for the *start* strain impeded the quantification of *aapA3*-Tr. For the G43A strain, the relative amount of *aapA3*-Tr mRNA in translating fractions was strongly reduced (approximately 4%, lanes 10 to 14, Figure 4B and Figure 4 – figure supplement 2E) compared to the T107C strain. This result confirmed that the single G43A mutation was sufficient to prevent ribosome binding, consequently impairing translation of the toxin. Remarkably, a similar result was observed for the A28C and A40T suppressor mutations. The A40T mutation had the strongest effect with only 7% of *aapA3*-Tr associated with translating ribosomes, compared to 19% for the A28C strain (Figure 4B and Figure 4 – figure supplement 2C, D). I*n vitro* translation assays (Figure 4 – figure supplement 1C) further confirmed that the A28C and A40T suppressor mutations, like the G43A mutant, act by reducing AapA3 translation efficiency.

Together, these results demonstrate that a single mutation within the 5’UTR either inside or outside the SD sequence, is able to overcome antitoxin absence by impeding toxin translation.

### The suppressor A28C and A40T mutations inhibit toxin translation by stabilizing a SD-sequestering hairpin

We next asked by which mechanism A28C and A40T mutations inhibit translation. Both substitutions lie in a single stranded region upstream of the minimal SD sequence (5’-AGGA-3’), which may be crucial for translation initiation (Figure 2). However, in both cases, a unique type of transversion mutation was selected (*i.e.*, A28T and A40C were not selected) suggesting that the mutations may act at the structure rather than at the primary sequence level. Indeed, secondary structure predictions with RNAfold algorithm (Gruber et al., 2008) suggested that both mutations could stabilize a local hairpin in which the SD is sequestered by an upstream aSD sequence (anti-SD sequence 1 (aSD1), Figure 5A). Indeed, while the A28C suppressor transversion extended this hairpin by one G-C base-pair, the A40T mutation created two additional A-U base-pairs. To investigate whether this stabilization was responsible for the translation inhibition effect, we tested whether the combination of A33T and A40T mutations (see Figure 5A), which is expected to destabilize the stem-loop stability, restored a toxic phenotype. Due to this potential toxicity, a strain containing an additional mutation in the AapA3 start codon was also generated (A33T/A40T/*start*, Figure 5B). Transformation assay was performed as previously described (Figure 3B). As expected, the suppressor A40T mutation was not toxic (Figure 5B). However, a 2 log-fold reduction in the number of Str^R^ transformants was observed with the A33T/A40T construct (Figure 5B). This effect disappeared when the toxin start codon was mutated (A33T/A40T/*start*, Figure 5B) demonstrating that the toxicity comes from the AapA3 peptide synthesis. This approach could not be used to study the A28C suppressor mutation since the non-compensatory mutation would lie within the SD sequence. Therefore, we tested the SD accessibility for all mutants (A40T, A28C, A40T/A33T) *in vitro* by performing an RNase H/oligonucleotide assay (Figure 5C, and Figure 5 – figure supplement 1). Compared to the WT and the A33T/A40T mutant, a reduced oligonucleotide accessibility was observed for both A28C and A40T *aapA3*-Tr RNAs, demonstrating that both mutations inhibit toxin expression by reducing SD accessibility.

**Figure 5.**
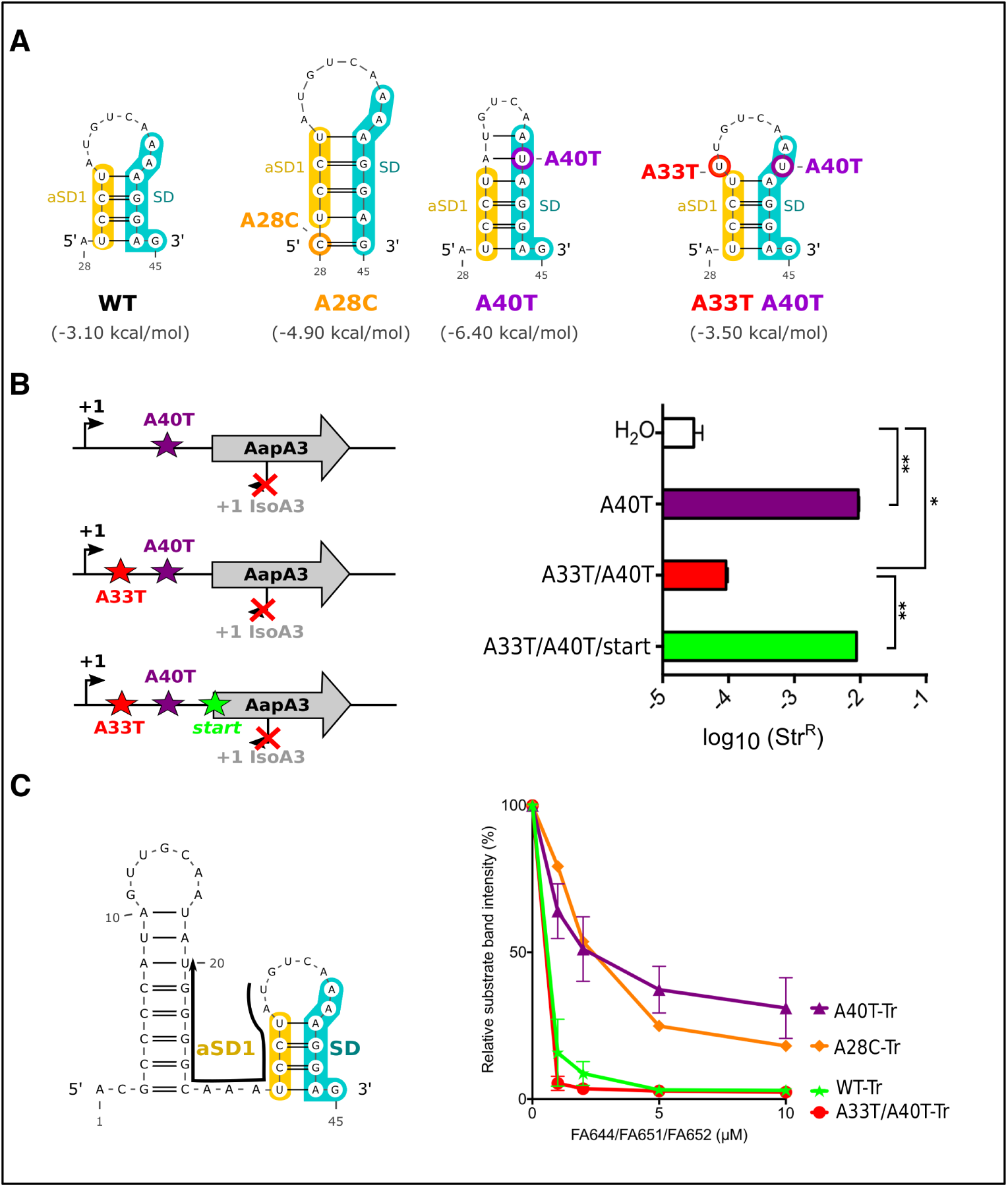
The A28C and A40T mutations suppress toxicity through SD-sequestration. **(A)** Secondary structures involving the SD sequence were predicted using the same software as in Figure 2. The A28C mutation (dark orange) generates an extra G-C base-pair; in the A40T mutant (purple), two additional A-U pairs stabilize the hairpin; the A40T antagonist mutation A33T is shown in red; the SD and the anti-SD (aSD1) sequences are shown in turquoise and in yellow, respectively. **(B)** (left) PCR constructs used to assess the SD sequestering structure by transformation assay. (right) For each transformation with the indicated PCR constructs, the number of Str^R^ obtained per total number of transformed cells was calculated and plotted on a log scale. Error bars represent s.d; *n=3* biological replicates. (\*\*\**P*<0.0001; \**P*=0.001 according to unpaired *t*-test). **(C)** Left: The position of the oligonucleotides (FA644 for WT and A40T; FA651 for A33T/A40T; and FA652 for A28C, see Table 4) used in the RNase H protection assay is indicated as a black arrow on the first 45 nucleotides of the *aapA3* mRNA. Right: 30 fmol of internally labeled WT and mutated *aapA3*-Tr transcripts were incubated with 0 to 100 pmoles of each specific DNA oligonucleotide and subjected to digestion by *E. coli* RNase H1. Digestion products were analyzed on an 8% PAA denaturing gel. Substrate consumption was quantified as relative substrate band intensity, 100% corresponding to the intensity obtained in absence of oligonucleotide. Error bars represent the s.d; *n=2* technical replicates.

Altogether these results demonstrate that both suppressor mutations are preventing translation initiation by stabilizing the SD sequestration within a local RNA hairpin instead of acting at the sequence level.

### A second SD-sequestering hairpin is embedded within the *aapA3* ORF

A synonymous substitution (T78C) that converts the Serine codon UCU into UCC was also selected in our suppressor selection. The presence of this mutation was intriguing since it is located 27 nt after the start codon and it does not affect the amino acid sequence. To understand its potential effect on toxin expression, we constructed a strain containing this mutation and the pIsoA3* mutation. Northern blot analysis showed that the T78C strain contains similar amounts of both *aapA3*-FL and -Tr mRNA isoforms than the other A28C and A40T suppressor strains (Figure 4 – figure supplement 1). We next tested the translatability of AapA3 *in vivo* by performing polysome fractionation coupled to Northern blot analysis. The percentage of *aapA3*-Tr found in the monosome and disome fractions of the T78C strain (Figure 6A and Figure 4 – figure supplement 2) was lower than that observed for the control T107C strain (34% vs 73%), but significantly higher than the one observed with the two A28C and A40T suppressors (Figure 4B). *In vitro* translation assays confirmed these results (Figure 4 – figure supplement 1), demonstrating that the T78C suppressor acts by inhibiting AapA3 translation.

**Figure 6.**
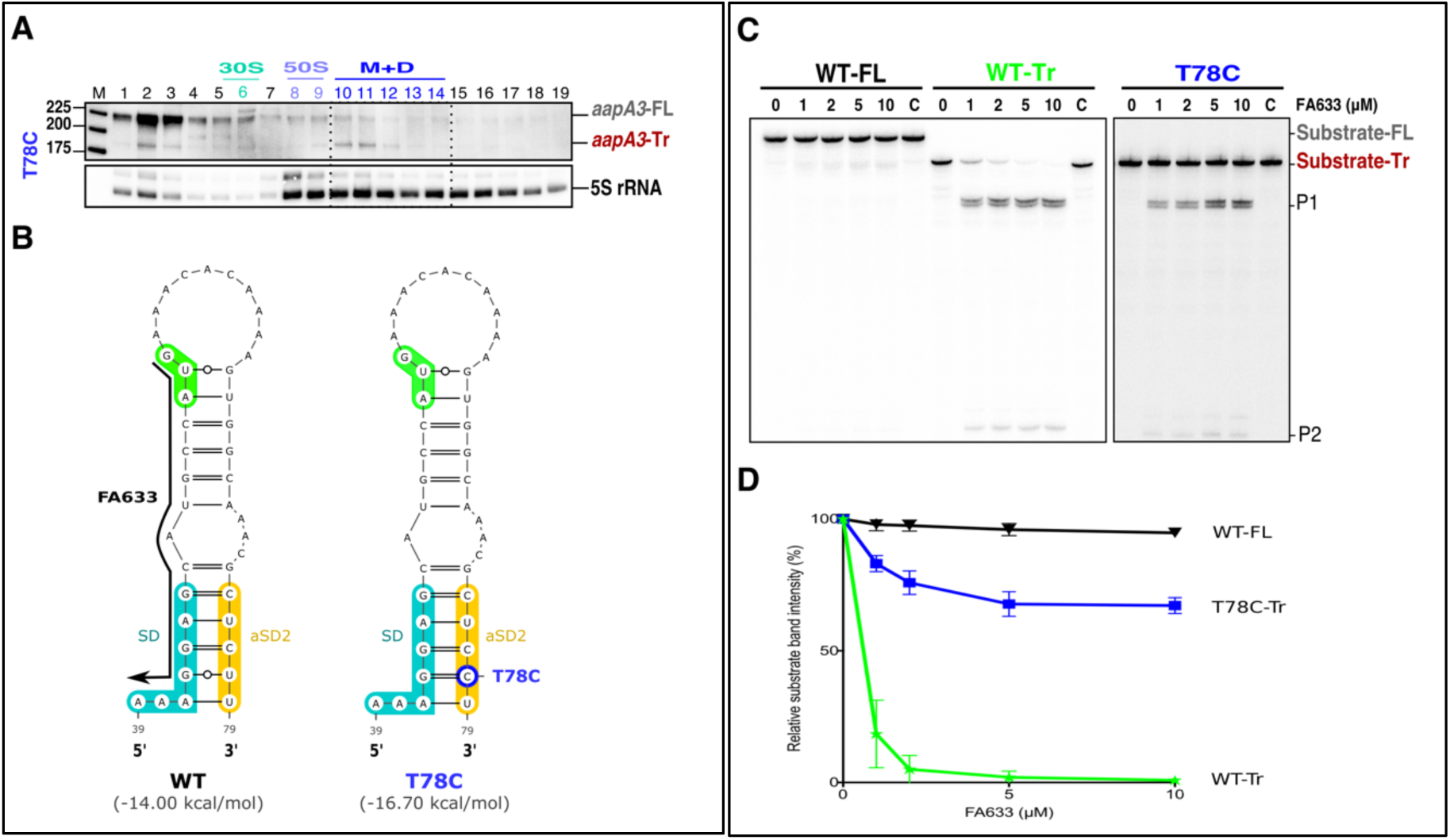
A synonymous mutation located within the toxin ORF inhibits *aapA3* mRNA translation via SD sequestration. **(A)** The cell lysate of T78C strain was subjected to ultracentrifugation through a sucrose gradient. RNA was analyzed as in Figure 4. The different transcripts *aapA3*-FL, *aapA3*-Tr, and 5S rRNA (loading control) are indicated. M+D, monosomes + disomes. **(B)** Prediction of the secondary structure involving the second aSD sequence. The black arrow represents the complementary sequence of the DNA oligonucleotide used in the assay. The T78C mutation is shown in dark blue, SD sequence in turquoise, anti-SD sequence in yellow and start codon in green. **(C)** A typical RNase H protection assays is shown. A total of 30 fmol of internally labeled *aapA3-*FL and *aapA3*-Tr RNA (WT or T78C) were incubated in presence of 0 to 100 pmoles of DNA oligonucleotide (FA633) and subjected to digestion by *E. coli* RNase H1. Lane C contains only the labeled substrate in absence of the enzyme. Two digestion products, P1 and P2 are indicated **(D)** Substrate consumption was quantified as the relative substrate band intensity and plotted as a function of DNA oligonucleotide concentration. Error bars represent the s.d; *n=2* technical replicates.

Secondary structure prediction revealed another putative SD-sequestering hairpin involving an aSD sequence (aSD2) embedded within the AapA3 ORF (Figure 6B). As for the two other A28C and A40T suppressor mutations, the T78C transition was expected to stabilize this hairpin by replacing a G-U by a G-C pair. To address the accessibility of this region, an RNase H protection assay was performed, using the FA633 oligonucleotide (Figure 6B). Remarkably, a strong reduction in SD accessibility was observed for the T78C RNA compared to the WT (Figure 6C and 6D). Thus, a single hydrogen bond is sufficient to stabilize the sequestration of the SD sequence and to suppress toxicity. Importantly, despite being located within the AapA3 coding region, the T78C suppressor acts at the mRNA folding level.

Sequence conservation analysis of the AapA3 coding region in 49 *H. pylori* strains (Figure 6 – figure supplement 1) revealed that the serine codon at position 9 is one of the most highly conserved codons of the peptide, indicating a crucial role of this sequence, likely in the sequestration of the SD sequence. Only the UM066 strain (highlighted in pink in Figure 6 – figure supplement 1) possesses a proline at this position, which probably abolishes peptide toxicity by disrupting the alpha-helix structure of the toxin (Masachis et al., 2018).

### Working Model

We have shown that mutations in aSD1 and aSD2 sequences suppress toxicity by stabilizing two mutually exclusive short hairpins that inhibit toxin translation (Figure 7). These stabilized hairpins are formed in the *aapA3*-Tr but not in the *aapA3*-FL (Figure 7 – figure supplement 1), explaining why they inhibit translation without affecting the stability of the full-length form. Interestingly, our FASTBAC-Seq approach revealed that, out of thirteen possible aSD sequences present in the *aapA3* mRNA (Figure 7 – figure supplement 1), only two could be mutated to suppress toxicity. Our results suggest that, in the WT context, these hairpins are not stable enough and are exclusively formed transiently during transcription. We propose that they act as functional metastable structures (MeSt1 and MeSt2, Figure 7), i.e. they form co-transcriptionally and sequentially to prevent premature toxin expression before a third aSD (aSD3) traps the *aapA3*-FL mRNA into a highly stable and translationally inert conformation (Figure 7).

**Figure 7.**
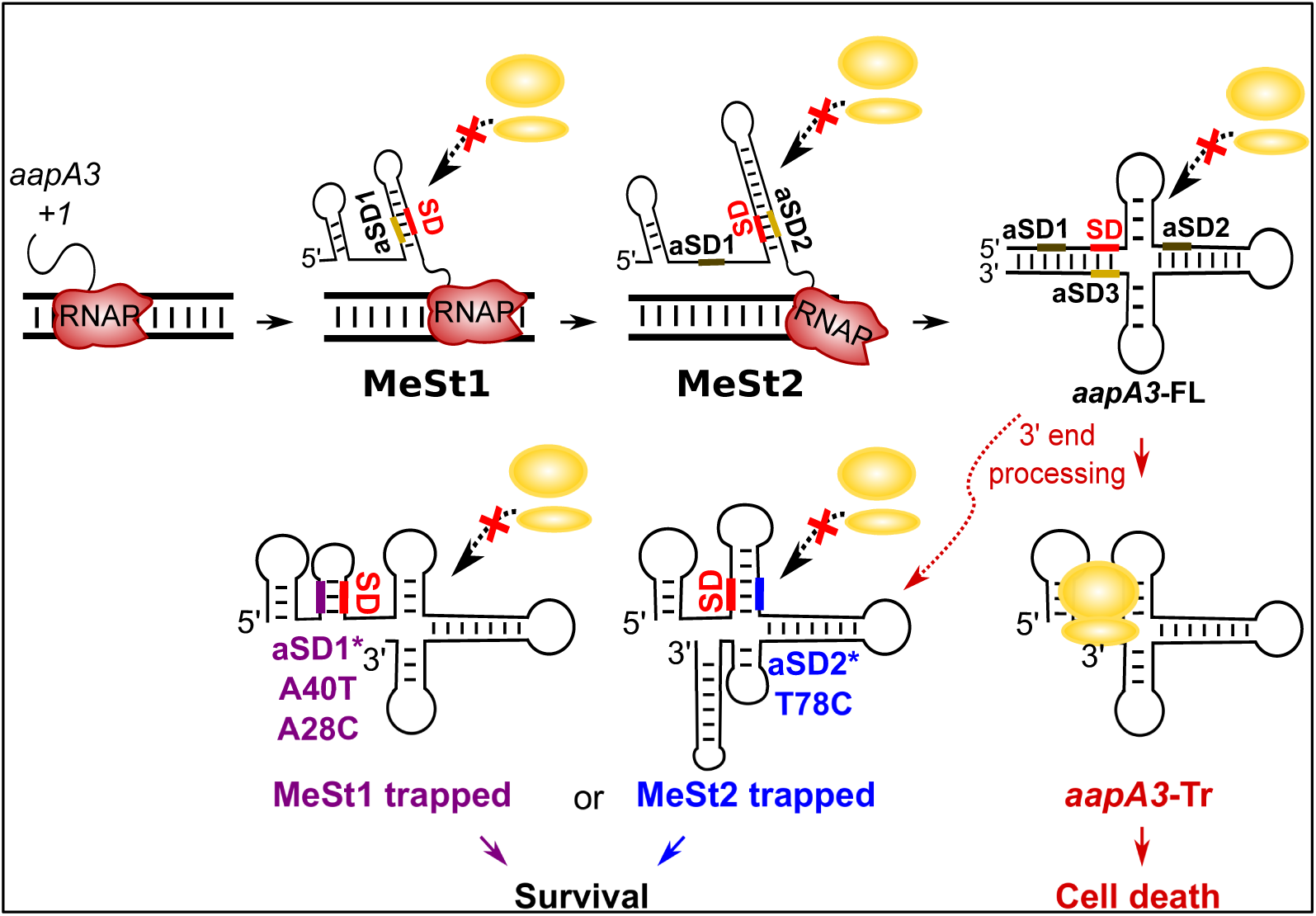
Working model of the *aapA3* co- and post-transcriptional regulation. Co-transcriptional folding of the *aapA3* mRNA leads to the generation of two successive SD-sequestering hairpins, which constitute metastable structures (MeSt) temporarily impeding ribosome access during transcription. The RNA polymerase is shown in red. Upon transcription termination, the *aapA3* full-length transcript (*aapA3*-FL) folds into a translationally inert conformation involving a 5’-3’-end long-distance interaction (LDI) in which the SD is sequestered by the aSD3 motif. A 3’-end nucleolytic truncation leads to the formation of *aapA3*-Tr, which is translationally active. In absence of IsoA3, its translation leads to cell death. We showed that the suppressor mutations A40T, A28C (purple) and T78C (blue) stabilize the metastable hairpins in the truncated isoform (involving the aSD1 and aSD2 sequences, respectively), leading to inhibition of *aapA3*-Tr translation. The suppressor strains can thus survive even in absence of IsoA3. The two successive MeSt structures act as thermodynamic traps to freeze the mRNA into translationally inert conformations.

## DISCUSSION

How bacteria modulate gene expression via RNA structure has been a fascinating topic for the last 30 years. This regulation is often achieved at the translation initiation step through the sequestration of the SD sequence in stable RNA hairpins that prevent ribosome binding to the RBS of the mRNA (Duval et al., 2013, 2015; Meyer, 2017b). Although recent advances using *in vivo* probing at the genome scale have confirmed that translation efficiency strongly correlates with the mRNA structure around the RBS (Mustoe et al., 2018), little is known about the influence of co-transcriptional folding on translation. Bacteria could, in principle, reduce or delay the translation of a specific mRNA by playing with its secondary structure while the mRNA is being made (Lai, Proctor, & Meyer, 2013; Zhu & Meyer, 2015). In this article, we identified two functional RNA hairpins within a type I toxin-encoding mRNA for which a tight control of translation is essential. We propose that these hairpins correspond to metastable structures that form sequentially and transiently to occlude the SD accessibility during mRNA synthesis.

### FASTBAC-Seq uses toxin lethality to identify suppressor mutations

To date, most studies on TA systems, including our previous work (Arnion et al., 2017), used artificial expression systems to characterize the effects of toxin expression. The use of such overexpression vectors is often a source of misinterpretations, as toxic proteins may not be found at such high concentrations under physiological conditions. To study the AapA3 toxin expression at the chromosomal level, we inactivated the endogenous IsoA3 antitoxin promoter as previously described for IsoA1 (Arnion et al., 2017). However, we were unable to obtain a viable strain without obtaining additional mutations in the toxin-encoding gene. Suppressor mutations have been also reported in *B. subtilis* for two chromosomally-encoded type I toxins with killer activity (*txpA*/RatA (Silvaggi et al., 2005) and *bsrG*/SR4 (Jahn et al., 2012)). Strikingly, a lethality at the chromosomal level has not been reported for chromosomally-encoded TA loci in Gram-negative bacteria. Indeed, the killer activity observed for the plasmid-encoded *hok*/Sok TA system was even believed to be not conserved for the chromosomally-encoded homologs (Pedersen & Gerdes, 1999). Interestingly, most of the *hok*/Sok homologs (*hokA*, *C* and *E*) in *E. coli* are inactivated by the presence of insertion elements located close to the toxin ORF (Pedersen & Gerdes, 1999). This observation, together with studies showing a differential expression of several TA systems in response to various stresses (*e.g.*, temperature shift, oxidative stress, starvation) (Harms et al., 2018), suggests that chromosomally-encoded TA systems may not be involved in a bactericidal activity, but rather, in a reversible growth arrest in response to a specific stress. Conversely, our results clearly demonstrate that, in line with the bactericidal activity observed for the overexpressed AapA1 toxin (Arnion et al., 2017), the chromosomal expression of the AapA3 toxin is constitutive and lethal in absence of the IsoA3 antitoxin. Consequently, we took advantage of this lethality to select suppressors and developed the FASTBAC-Seq method to rapidly identify hundreds of intragenic suppressor mutations with nucleotide resolution (Masachis et al., 2018).

### A single-nucleotide substitution is sufficient to impede toxin translation

The FASTBAC-Seq method revealed a wide range of unanticipated *cis*-encoded toxicity determinants, affecting either the toxic activity of the protein (described in (Masachis et al., 2018), or its expression (this study). Among the mutations affecting the toxin mRNA expression, we identified five single-nucleotide substitutions able to inhibit the translation of AapA3 mRNA without affecting its stability. Three of them were located in the SD sequence. The most highly enriched mutations substituted the guanines at positions 42 and 43 by either an adenine, a cytosine, or an uridine. This revealed 5’-AGG-3’ and 5’–GGA-3’ as the minimal functional SD motifs allowing AapA3 translation, in agreement with the previously identified *H. pylori* SD consensus sequence (5’-AAGGA-3’) (C. M. Sharma et al., 2010). The third mutation (A40T) was much less enriched, and remarkably, only the transversion from an adenine to a thymine was selected. Another transversion mutation (A28C) was selected 14 nt upstream of the SD sequence. The fact that only transversion mutations were selected at these two positions indicated that the nature of the substituted nucleotide was important, suggesting that they may not directly act at the sequence, but rather at the structure level. Indeed, our results demonstrated that the A28C and A40T mutations create, respectively, one or two additional base-pair(s) within a stem-loop structure formed by the pairing between the SD sequence and an upstream complementary aSD sequence (aSD1, 5’-UCCU-3’). Destabilizing the A40T mutated stem by mutating the complementary nucleotide (A33T) restored toxicity, clearly showing that the A40T mutation, despite being located within the SD sequence, acts at the mRNA structural level and not at the sequence level.

Interestingly, the T78C mutation revealed the existence of a second aSD sequence (aSD2) located downstream the SD sequence, within the toxin coding region. This synonymous substitution (UC<u>U</u>→UC<u>C</u>, Ser codon at position 9) creates a perfect aSD sequence (5’-CUCCU-3’). Although this mutation could potentially create a rare codon reducing toxin translation efficiency, we did not favor this hypothesis since the less frequently used Ser codon in *H. pylori* is UCG (Atherton, Sharp, & Lafay, 2000). Interestingly, synonymous mutations close to the translation initiation region (TIR) have also been shown to influence gene expression by modulating the stability of mRNA folding rather than by acting at the codon usage level (Kudla, Murray, Tollervey, & Plotkin, 2009). In addition, a strong codon bias has also been observed within the first 15 codons, which avoids tight mRNA structure close to the TIR region (Bentele, Saffert, Rauscher, Ignatova, & Bluthgen, 2014; Bhattacharyya et al., 2018). Here, we showed that despite the presence of up to thirteen CU-rich sequences in the AapA3 mRNA, only mutations in the sequences closest to the SD could be selected, reflecting a distance-dependence of these translation regulatory elements. A similar aSD sequence (5’-UCCU-3’) has been identified in the coding sequence of the *gnd* gene in *E. coli* (Carter-Muenchau & Wolf, 1989). Interestingly, displacing this aSD sequence from its natural position (codon 66) to a more proximal position (codon 13) greatly increased its capacity to inhibit translation.

### Suppressor mutations reveal functional metastable structures acting co-transcriptionally to impede premature toxin translation

The three mutations studied here (A28C, A40T and T78C) act post-transcriptionally after the 3’ end processing by stabilizing SD-sequestering hairpin structures. Importantly, these suppressor mutations do not interfere with the folding pathway of the full-length mRNA, neither affecting its transcription, stability, nor its 3’-end maturation, indicating that they exclusively act on the active, truncated, AapA3 mRNA form (Figure 7). Interestingly, these local hairpins were previously predicted to form during the co-transcriptional folding pathway of several AapA mRNAs (Arnion et al., 2017). Now, our FASTBAC-seq approach reveals that these structures are functional, *i.e.* they transiently form during transcription to prevent toxin translation before the FL is made. Indeed, stabilizing these hairpins inhibits translation of the *aapA3*-Tr. This temporal control of gene expression is achieved through the sequential formation of two RNA hairpin structures that mask the SD sequence via CU-rich elements. In the full-length mRNA, these structures are replaced by a more stable one involving an LDI between both ends of the transcript. Similar to the *hok* mRNA, this final mRNA structure is so stable that its translational activation requires a 3’-end processing removing the aSD3 sequence element. The highly stable structure of the *aapA3*-FL mRNA is also similar to the cloverleaf-like structure present in the 5’ UTR of the MS2 coliphage maturation gene (Groeneveld, Thimon, & van Duin, 1995). Interestingly, in this case, it may take up to several minutes for the mRNA to be synthesized and properly folded (van Meerten, Girard, & van Duin, 2001), explaining the need of functional transient structural intermediates preventing premature gene expression.

The selection of three stabilizing mutations suggests that the thermodynamic stability of such SD-sequestering stem-loops in the WT context is not sufficient to inhibit the translation of the active AapA3 mRNA form. Instead, our results suggest that in the WT situation, these SD-sequestering hairpins (MeSt1 and MeSt2, Figure 7) are only transiently formed to co-transcriptionally impede the premature toxin translation. This transient character is essential to ensure the proper transcription termination and folding of the full-length mRNA, and it is achieved by hierarchically increasing thermodynamic stabilities (Figure 7 – figure supplement 2). Importantly, the suppressor mutations do not provide enough stabilization to impede the formation of the next most stable structure. Indeed, the A40T mutated MeSt1 has an energy of −21.10 kcal/mol, while that of the WT MeSt2 is −29.30 kcal/mol (Figure 7 – figure supplement 2). This may explain why the SD-sequestering mutations do not interfere with its co-transcriptional folding pathway and why the last SD-aSD3 is finally formed in the mutants.

The importance of metastable RNA structures in the AapA3 mRNA is attested by the strict conservation of the UCU Serine codon at position 9. As our results have shown, a synonymous UCC codon at this position (T78C mutation) would inhibit the AapA3 toxin expression, rendering the TA locus non-functional and probably promoting its rapid loss. Our study represents the first *in vivo* evidence of the existence of sequential RNA metastable structures that avoid, directly but transiently, the co-transcriptional translation of a toxin-encoding mRNA.

The formation of metastable structures has been reported in several RNA-mediated regulatory pathways, including viral RNA replication (Repsilber et al., 1999), RNA catalysis (Pan & Woodson, 1998), RNA editing (Linnstaedt, Kasprzak, Shapiro, & Casey, 2006), and ribosome biogenesis (I. M. Sharma et al., 2018). They are usually described as folding intermediates that work in a hierarchical manner to help an RNA molecule reaching its functional conformation (*i.e.*, most thermodynamically stable conformation). A nice example of such metastable structures has been reported for the regulation of the *hok*/Sok type I TA system in *E. coli*. In this pioneering work, they showed that the formation a metastable hairpin ensures the proper folding of the Hok mRNA into a translationally inert conformation (Møller-Jensen, Franch, & Gerdes, 2001; Nagel, Gultyaev, Gerdes, & Pleij, 1999). Although this metastable hairpin is located at the 5’ end of the mRNA, it does not directly mask the SD sequence. Instead, it favors a conformation in which the SD is sequestered by a downstream anti-SD. Other metastable structures are directly involved in the activation or inhibition of gene expression (Zhu & Meyer, 2015), as examplified by the structures reported in the Trp operon leader, the SAM riboswitch and the 5’ UTR of the MS2 phage (Zhu & Meyer, 2015). The metastable structures of *aapA3* are more reminiscent of the latter example (van Meerten et al., 2001), except that in the case of MS2, the transient structure allows translation to occur before the cloverleaf-like structure is formed. Nevertheless, in both cases, a functional transient RNA structure exerts a temporal control of translation, either negatively or positively.

### Conclusion

Although the coupling between transcription and translation in bacteria plays important roles in gene expression (Kriner, Sevostyanova, & Groisman, 2016), it can be harmful in the case of toxin-encoded mRNAs. Thus, the metastable RNA structures identified in the present study are essential to uncouple transcription and translation processes and allow the presence of type I TA systems on bacterial chromosomes. Although transient RNA structures can be predicted *in silico* (Meyer, 2017), their *in vivo* characterization remains challenging. Several high-resolution methods have been recently reported for analyzing the co-transcriptional folding of regulatory RNAs, both *in vitro* (Uhm, Kang, Ha, Kang, & Hohng, 2018; Watters, Strobel, Yu, Lis, & Lucks, 2016) and *in vivo* (Incarnato et al., 2017). These complementary techniques may be useful to analyze the formation of these metastable hairpins in real-time.

## MATERIALS AND METHODS

### Bacterial Strains, Plasmids and Growth conditions

The *H. pylori* strain used in this study is the 26695 reference strain (Tomb et al., 1997). Strains were grown on Columbia agar plates supplemented with 7% horse blood and Dent selective supplement (Oxoid, Basingstoke, UK) for 24 to 48 h depending on the strain. Liquid cultures were performed in Brain-Heart Infusion (BHI) medium (Oxoid) supplemented with 10% fetal bovine serum (FBS) and Dent. *H. pylori* plates and liquid cultures were incubated at 37°C under microaerobic conditions (10% CO_2_, 6% O_2;_ 84% N_2_) using an Anoxomat (MART microbiology) atmosphere generator. Plasmids used for cloning were amplified in *Escherichia coli* TOP10 strain, which was grown in Luria-Bertani (LB) media, supplemented either with kanamycin (50 μg.mL^−1^), chloramphenicol (30 μg.mL^−1^) or ampicillin (100 μg.mL^−1^). For *H. pylori* mutant selection and culture, antibiotics were used at the following final concentrations: 20 μg.mL^−1^ kanamycine (Sigma), 8 µg.mL^−1^ chloramphenicol (Sigma), 10 μg.mL^−1^ streptomycin and 10 μg.mL^−1^ erythromycin.

### Molecular techniques

Molecular biology experiments were performed according to standard procedures and the supplier recommendations. High Purity Plasmid Miniprep Kit (Neo Biotech) and Quick Bacteria Genomic DNA extraction Kit (Neo Biotech) were used for plasmid preparations and *H. pylori* genomic DNA extractions, respectively. PCR were performed either with Dream Taq DNA polymerase (Thermo Fisher Scientific), or with Phusion High-Fidelity Hot Start DNA polymerase (Thermo Fisher Scientific) when the product required high-fidelity polymerase. Site-directed mutagenesis PCR was performed with the PfuUltra High-Fidelity DNA Polymerase (Agilent). All oligonucleotides used in this study are shown in Table 4.

### RNA extraction

For RNA extraction, bacterial growth was stopped at the desired OD_600nm_ by adding 650 μl cold Stop Solution (95% ethanol, 5% phenol pH 4.5) to 5 ml of culture, which was placed on ice. Cells were then centrifuged for 10 min at 3,500 rpm and 4°C, and the pellets were stored at −80°C. Cell pellets were resuspended in 600 μl Lysis Solution (20 mM NaAc pH 5.2, 0.5% SDS, 1 mM EDTA) and added to 600 µl hot phenol pH 5.2. After incubation for 10 min at 65°C, the mixture was then centrifuged for 10 min at 13,000 rpm and room temperature. The aqueous phase was next transferred to a phase-locked gel tube (Eppendorf) with an equal volume of chloroform and centrifuged for 10 min at 13,000 rpm and room temperature. Total RNA was then precipitated from the aqueous phase by adding 2.5 volumes of ethanol 100% and 1/10 volume of 3 M NaAc pH 5.2. After centrifugation for 30 min at 13,000 rpm and 4°C, the supernatant was discarded and the pellet was washed with 75% ethanol. Finally, the supernatant was discarded and the RNA pellet air-dried and resuspended in H_2_O. For RNA half-life determinations, rifampicin (Sigma, prepared at 34 mg.ml^−1^ in methanol) was added to the culture at a final concentration of 80 µg.ml^−1^ and cells were harvested at the desired time points. A culture where rifampicin was replaced by the same volume of methanol served as a non-treated control.

### Northern Blot

For Northern blot analysis, 1 to 10 μg RNA were separated on an 8% polyacrylamide (PAA), 7M urea, 1X Tris Borate EDTA (TBE) gel. RNA was transferred to a nylon membrane (Hybond^TM^-N, GE Healthcare Life Science) by electroblotting in TBE 1X at 8V and 4°C overnight. Then, RNA was cross-linked to the membrane by UV irradiation (302 nm) for 2 min in a UV-crosslinker and hybridized with 5’-labeled (γ^32^P) oligodeoxynucleotides in a modified Church Buffer (1 mM EDTA, 0.5 M NaPO_4_ pH 7.2, 7% SDS) overnight at 42°C. Membranes were washed two times 5 minutes in 2X SSC, 0.1% SDS, and revealed using a Pharos FX phosphorimager (Biorad). For riboprobes, a DNA template containing a T7 promoter sequence was amplified by PCR from *H. pylori* 26695 genomic DNA as template. *In vitro* transcription was performed as described in the MaxiScript T7 Transcription Kit (Ambion) in the presence of 50 µCi of ^32^P-α-UTP and 1 mM cold UTP and purified on a Sephadex G25 column (GE Healthcare). Hybridization was performed in the modified Church Buffer at 65°C and the membrane was washed two times 5 min in 2X SSC, 0.1% SDS at 65°C. For the detection of *aapA3* mRNA species the ^32^P-labelled primer FD38 was used. To detect the *aapA3* mutants sequestering the SD region (where the primer FD38 binds), a riboprobe corresponding to the 5’ UTR of the mRNA was transcribed from a PCR fragment containing the T7 promoter and amplified with the FA170/FA11 primer pair. IsoA3 RNA was detected with a riboprobe corresponding to the *aapA3*-Tr RNA species transcribed from a PCR fragment containing the T7 promoter and amplified with the primer pair FA170/FA173.

### *In vitro* transcription and translation assays

For *in vitro* synthesis of the *aapA3* and IsoA3 RNAs, DNA templates were amplified from *H. pylori* 26695 genomic DNA using primer pairs: FA170/FA175 (*aapA3*-FL), FA170/FA173 (*aapA3-*Tr), FD11/FD17 (IsoA3), each forward primer carrying a T7 promoter sequence (see primer list, Table 4). *In vitro* transcription was carried out using the MEGAscript® T7 Transcription Kit (Ambion #AM1334) according to the manufacturer’s protocol. After phenol:chloroform extraction followed by isopropanol precipitation, the RNA samples were desalted by gel filtration using a Sephadex G-25 (GE Healthcare) column. For *in vitro* translation of the *aapA3*-FL and *aapA3-*Tr mRNAs, 0.5 µg of RNA was added to the *E. coli* S30 Extract System for Linear Templates Kit (Promega #L1030) as previously described (C. M. Sharma et al., 2010).

### *In vitro* structure probing

20 pmol of both *aapA3-FL* and *aapA3-Tr* transcripts were dephosphorylated with 10 U of calf alkaline phosphatase (CIP) at 37°C for 1 h. RNA was isolated by phenol extraction and precipitated overnight at −20°C in the presence of 30:1 ethanol: 0.3M NaOAc pH 5.2 and 20 µg GlycoBlue^TM^. The dephosphorylated RNA was then 5’ end-labelled with 10 pmol ^32^P-γ-ATP using the T4 polynucleotide kinase (PNK) for 30 min at 37 ° C. Unincorporated nucleotides were removed using a MicroSpin^TM^ G-25 column and labelled RNA was purified on an 8% PAA gel containing 7 M urea and 1X TBE. Upon visualization of the labelled RNA, the band corresponding to the RNA species of interest was cut from the gel and eluted overnight at 4°C under shaking in 750 µl RNA elution buffer (0.3M NH_4_Ac, 0.1% SDS, 1mM EDTA). RNA was extracted by Phenol/Chloroform/Isoamyl alcohol (25:24:1 v/v), and precipitated by ethanol (2.5V), pellets were washed and resuspended in 50 µl H_2_O and stored at −20°C.

Before use, each *in vitro* transcribed RNA was denatured by incubation at 90°C for 2 min in the absence of magnesium and salt, then chilled on ice for 1 min, followed by a renaturation step at room temperature for 15 min in 1X Structure Buffer (10 mM Tris-HCl pH 7.0, 10 mM MgCl_2_, 100 mM KCl). Structure probing analyses were performed as described previously (Darfeuille, Unoson, Vogel, & Wagner, 2007; C. M. Sharma et al., 2010; C. M. Sharma, Darfeuille, Plantinga, & Vogel, 2007), using 0.1 pmol of 5’ end-labeled RNA. To determine the secondary structure of RNA, 1 µl RNase T1 (0.01 U.μl-1; Ambion) was added to the labeled RNA and incubated in 1X Sequencing Buffer (20 mM Sodium Citrate, pH 5.0, 1 mM EDTA, 7M Urea) for 5 min at 37°C. Lead acetate (5 mM final concentration) digestions of both *aapA3*-Tr and *aapA3*-FL were done in the absence or in the presence of 2-10-fold excess of cold IsoA3 RNA. All reactions were stopped by adding 10 µl of 2X Loading Buffer (95% formamide, 18 mM EDTA, Xylene Blue and Bromophenol Blue. Cleaved fragments were then analyzed on an 8% denaturing PAA gel containing 7M urea and 1X TBE. Gels were dried for 45 min at 80°C, and revealed using a Pharos FX phosphorimager (Biorad).

### RNase H1/oligonucleotide accessibility assay

Internally-labelled transcripts were *in vitro*-transcribed using the MAXIscript® T7 Transcription Kit (Ambion #AM1312) in presence of 2.2 μM α-^32^P-UTP according to the manufacturer’s protocol. Labeled RNA was purified on an 8% PAA gel containing 7 M urea and 1X TBE, eluted overnight at 4°C under shaking in 750 µl elution buffer (0.1 M NaOAc pH 5.2, 0.1% SDS). RNA was desalted and concentrated by ethanol precipitation, pellets were resuspended in 100 µl H_2_O. Approximately 30 fmol of RNA were used for RNase H/oligonucleotide accessibility assays. Before use, each *in vitro*-transcribed RNA and DNA oligonucleotides were denatured as described for structure probing. Next, DNA oligonucleotides complementary to the region around the SD sequence (FA633 for WT and T78C (aSD2) mRNA; FA644 for A40T mRNA (aSD1); FA651 for the double mutant A33T/A40T mRNA (aSD1); and FA652 for A28C mRNA) were added to a final concentration of 0 to 10 μM. Reactions were adjusted to a final volume of 10 μl with H_2_O and incubated for 30 min at 30°C in the presence or absence (control) of 0.25 U *E. coli* RNase H1 (Ambion #AM2293). Reactions were then stopped by addition of 10 μl of 2X Loading Buffer (95% formamide, 18 mM EDTA, Xylene Blue and Bromophenol Blue). Cleaved fragments were analyzed on an 8% denaturing PAA gel containing 7M urea and 1X TBE. Gels were dried 45 min at 80°C, and revealed using a Pharos FX phosphorimager (Biorad).

### *H. pylori* chromosomal manipulation techniques

All mutant *H. pylori* strains listed in Table 1 were generated by chromosomal homologous recombination of PCR-generated constructs, introduced by natural transformation, as previously described (Masachis et al., 2018). In all cases, constructs contained ≈ 400 nt of the up- and downstream chromosome regions of the target gene, flanking the DNA fragment to be introduced (*i.e.*, antibiotic resistance marker to generate deletions or a WT copy of the target gene for complementation). DNA fragments of interest were previously cloned in *E. coli* vectors to avoid *H. pylori* WT genomic DNA (gDNA) contamination (see ‘*aapA3*/IsoA3 locus sub-cloning in *E. coli*’ section below). Constructs were generated by PCR assembly of PCR products amplified from the plasmids listed in Table 2 with the oligonucleotides shown in Table 4. Prior to transformation, *H. pylori* strains (number of cells corresponding to 1 OD_600nm_) were grown on non-selective CAB plates. After 4 hours incubation at 37°C under microaerobic conditions, 1 μg of PCR assembly product was added to the cells and plates were incubated for another 16 hours. Transformed cells were then selected on plates supplemented with the appropriate antibiotics and incubated for 4-6 days until isolated colonies appeared. Genomic DNA from transformants was purified using the Quick Bacteria Genomic DNA extraction Kit and subjected to PCR and Sanger sequencing for mutant validation.

**Table 1.**
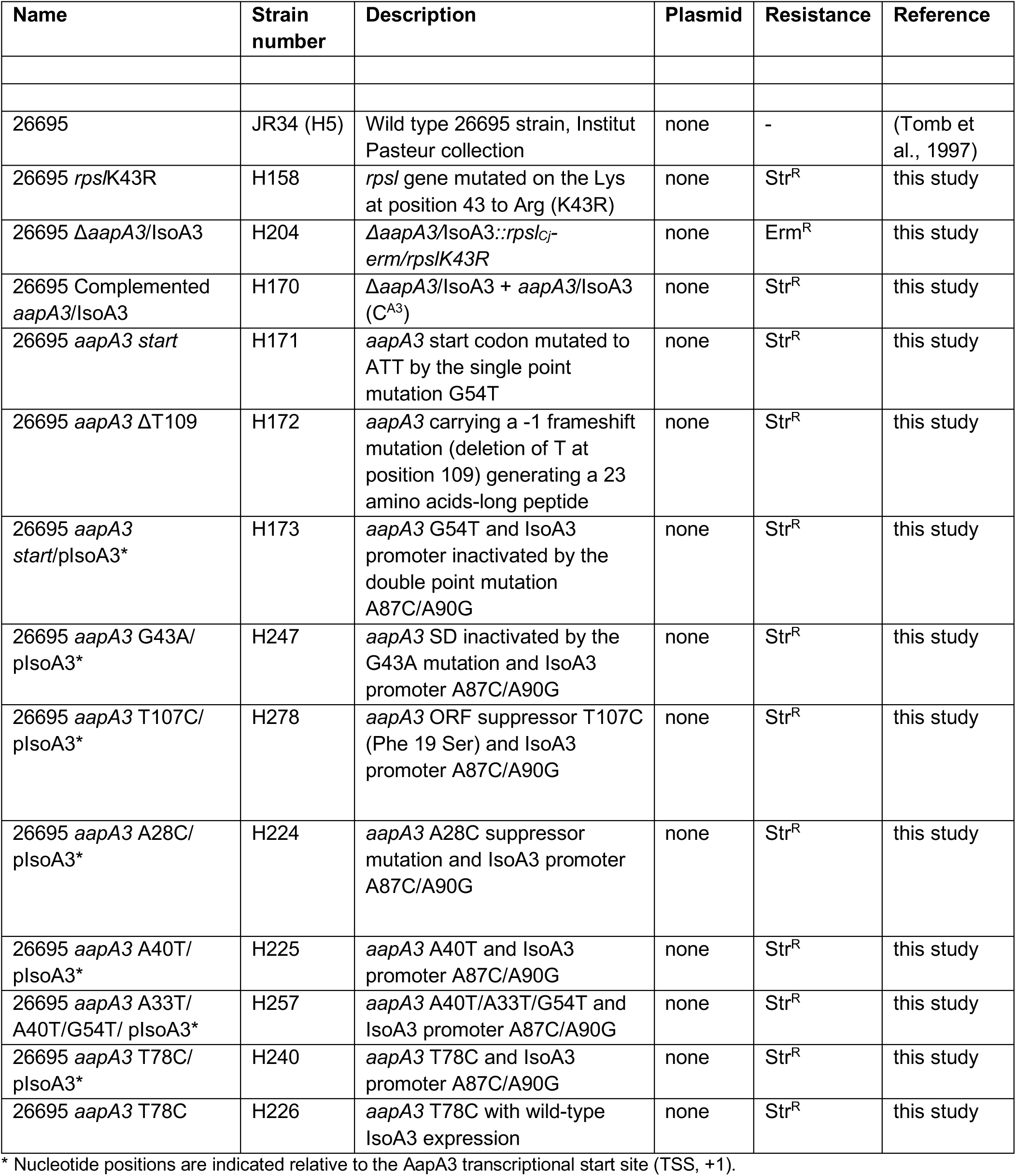
*Helicobacter pylori* strains used in this work.

**Table 2.** Plasmids used in this work.

| Name | Description | Origin/<br>Marker | Reference |
| --- | --- | --- | --- |
| pSP60 -2 | pSP60 carrying the counter selection cassette *rpsL-erm* | pSC101*/<br>Amp<sup>R</sup> | (Dailidiene, D. *et al.*, 2006)<br>(Pernitzsch *et al.*, 2014) |
| pA3-Up WT | pGEM-T carrying the upstream fragment of the *aapA3/IsoA3* locus amplified with the FA406/FA386 primer pair | ColE1/<br>Amp<sup>R</sup> | this study |
| pA3-Down WT | pGEM-T carrying the downstream fragment of the *aapA3/IsoA3* locus amplified with the FA409/FA387 primer pair | ColE1/<br>Amp<sup>R</sup> | this study |
| pA3-Down pIsoA3* | pGEM-T carrying the downstream fragment of the *aapA3/IsoA3* locus containing IsoA3 -10 box inactivated (A87C/A90G) | ColE1/<br>Amp<sup>R</sup> | this study |
| pA3-Up *start* | pGEM-T carrying the upstream fragment of the *aapA3/IsoA3* locus containing the AapA3 start codon mutation G54T | ColE1/<br>Amp<sup>R</sup> | this study |
| pA3-Up A28C | pGEM-T carrying the upstream fragment of the *aapA3/IsoA3* locus containing the suppressor A28C | ColE1/<br>Amp<sup>R</sup> | this study |
| pA3-Up A33T | pGEM-T carrying the upstream fragment of the *aapA3/IsoA3* locus containing the A33T mutation | ColE1/<br>Amp<sup>R</sup> | this study |
| pA3-Up A40T | pGEM-T carrying the upstream fragment of the *aapA3/IsoA3* locus containing the suppressor A40T | ColE1/<br>Amp<sup>R</sup> | this study |
| pA3-Up A40T/A33T | pGEM-T carrying the upstream fragment of the *aapA3/IsoA3* locus containing the A40T and the compensatory mutation A33T | ColE1/<br>Amp<sup>R</sup> | this study |
| pA3-Up T78C | pGEM-T carrying the upstream fragment of the *aapA3/IsoA3* locus containing the suppressor T78C | ColE1/<br>Amp<sup>R</sup> | this study |
| pA3-Down T78C | pGEM-T carrying the downstream fragment of the *aapA3/IsoA3* locus containing the suppressor T78C | ColE1/<br>Amp<sup>R</sup> | this study |
| pA3-Down T78C/ pIsoA3* | pGEM-T carrying the downstream fragment of the *aapA3/IsoA3* locus containing the suppressor T78C and IsoA3 -10 box inactivated (A87C/A90G) | ColE1/<br>Amp<sup>R</sup> | this study |
| pA3-Up G43A | pGEM-T carrying the upstream fragment of the *aapA3/IsoA3* locus containing the SD suppressor G43A | ColE1/<br>Amp<sup>R</sup> | this study |

\* Nucleotide positions are indicated relative to the AapA3 transcriptional start site (TSS, +1).

**Table 3.**
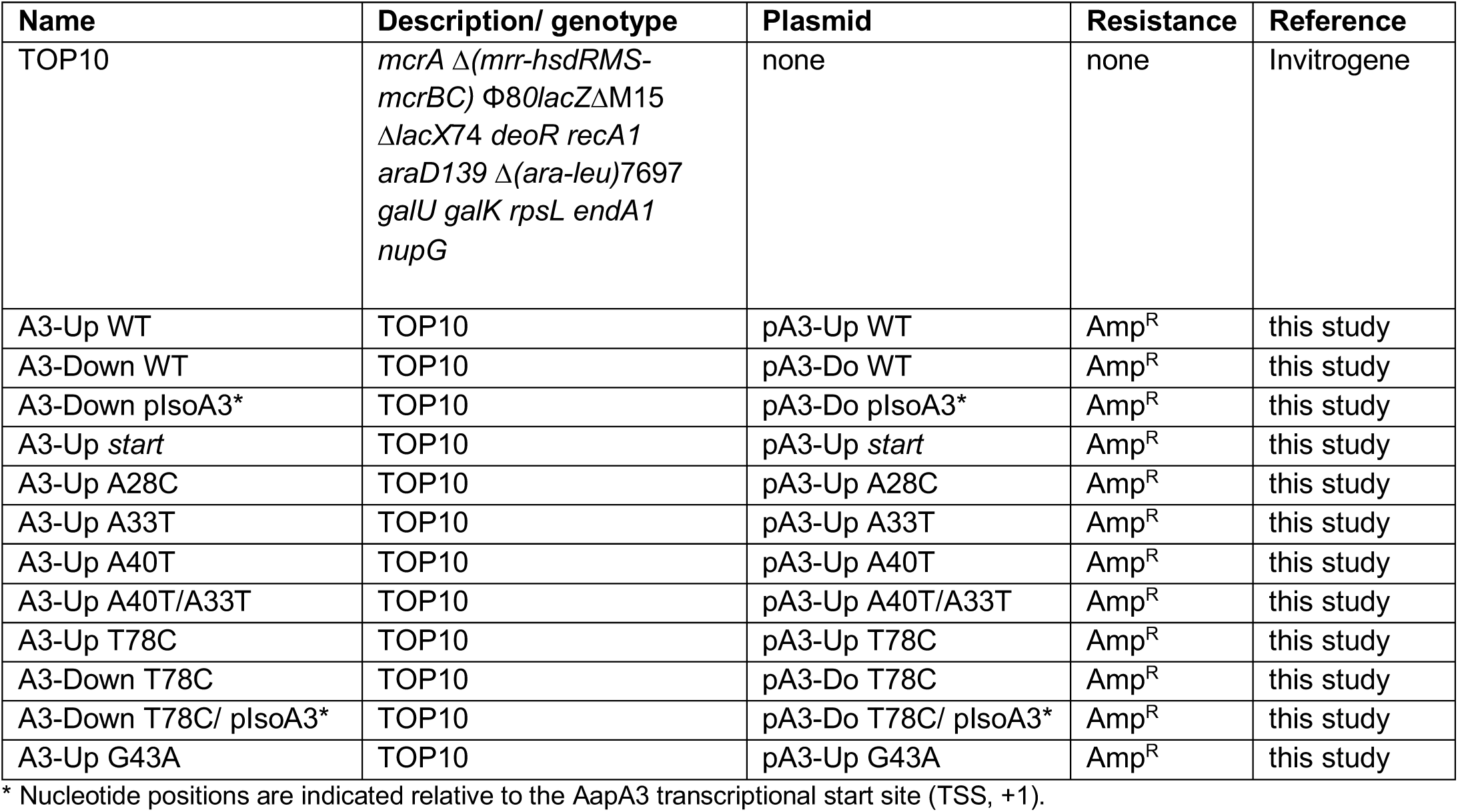
*Escherichia coli* strains used in this work.

**Table 4.**
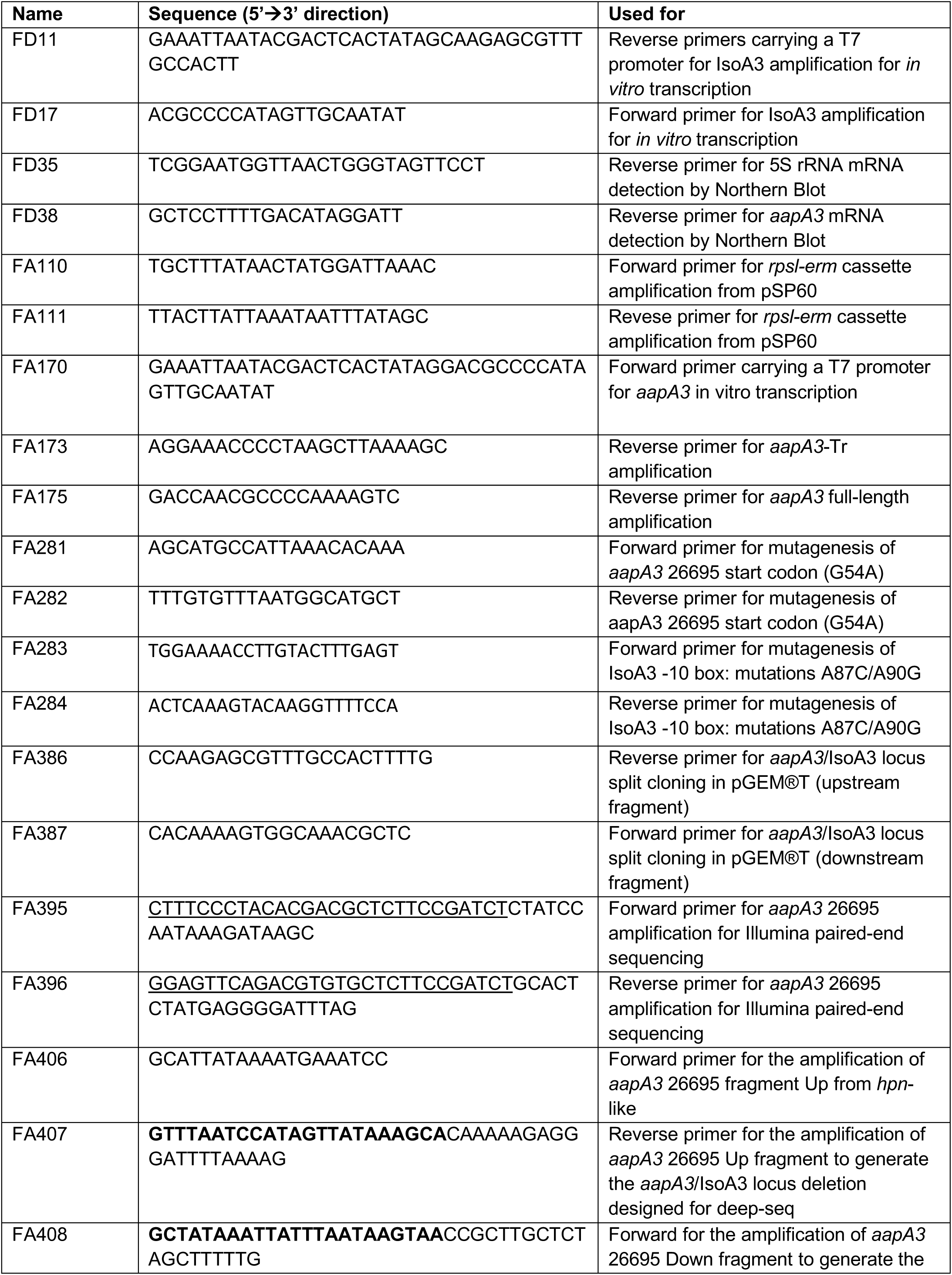
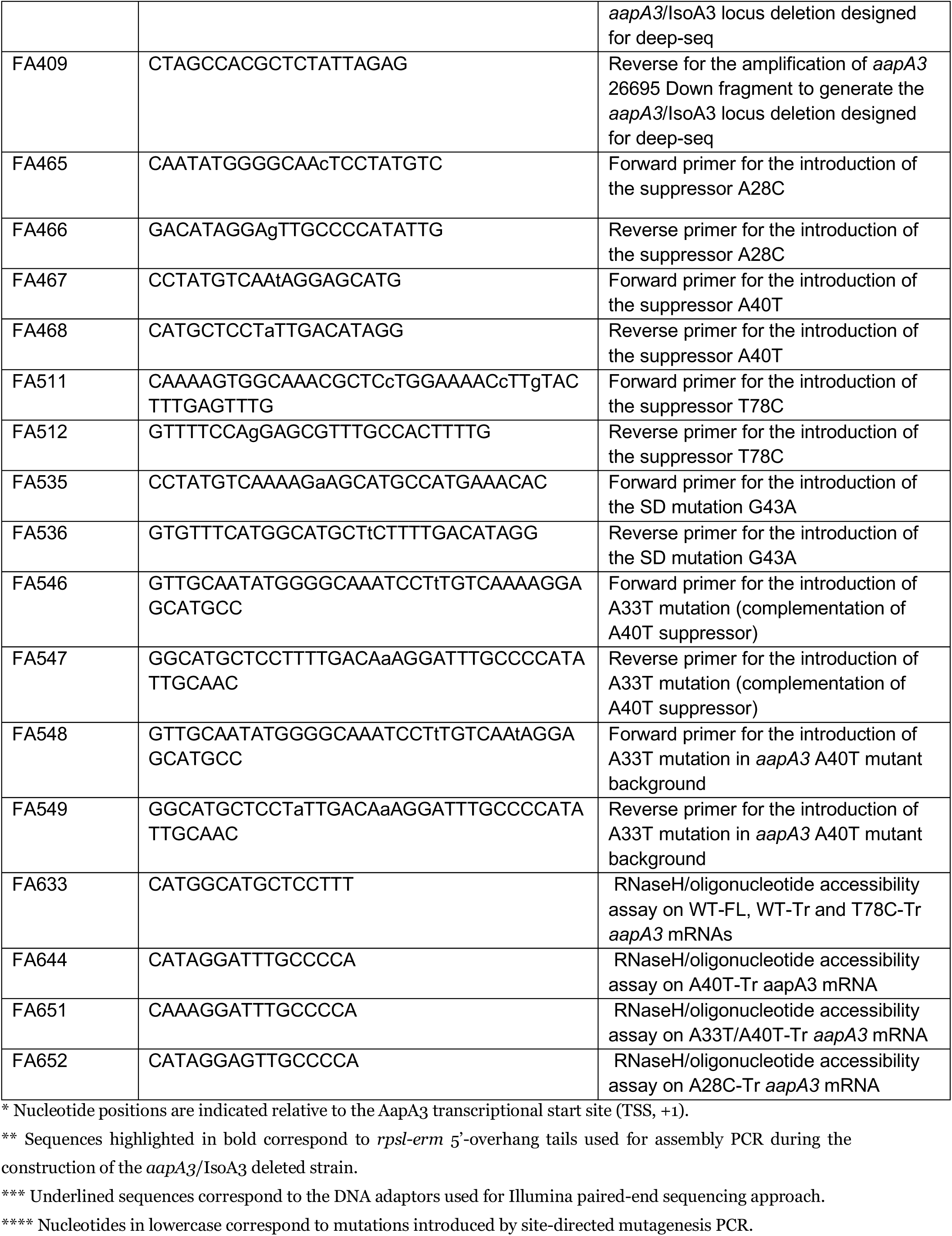
Oligonucleotides used in this work.

### Deletion of the *aapA3*/IsoA3 locus using the *rpsl_Cj_-erm* counterselection marker

The counterselection cassette *rpsl_Cj_-erm* was used to generate an *H. pylori* 26695 strain deleted for the *aapA3*/IsoA3 locus following the protocol described in (Masachis et al., 2018). First, the 26695 *H. pylori* strain used in this study was modified in order to become resistant to streptomycin. To this end we introduced by homologous recombination a mutation (K43R) in the rpsL gene coding for the small S12 ribosomal protein (Masachis et al., 2018). Then, up- and downstream fragments to the locus were amplified with the primer pairs FA406/FA407 and FA408/FA409. These flanking regions (415 and 418 nt-long, respectively) allow chromosomal homologous recombination to occur. The internal primers (FA407 and FA408) were used to introduce a 3’- and 5’- *rpsl_Cj_-erm* cassette homology tail, respectively, to allow subsequent PCR assembly. The *rpsl_Cj_-erm* cassette was amplified from the pSP60-2 plasmid (Table 2) using the primer pair FA110/FA111. Then, the up- and downstream fragments were assembled with the *rpsl_Cj_-erm* cassette by PCR assembly using the external primers (FA406/FA409) (see Figure 1—figure supplement 1 and primer list in Table 4). This construct (1294 nt-long) was used to perform natural *H. pylori* transformation by homologous recombination, as previously described (Bury-Moné, Skouloubris, Labigne, & De Reuse, 2001). This process generated the strain that will serve as recipient in all our successive transformation experiments, Δ*aapA3*/IsoA3::*rpsl_Cj_-erm*/K43R (Table 1).

### *aapA3*/IsoA3 locus sub-cloning in *E. coli*

Because *H. pylori* has a highly active homologous recombination machinery, a cloning step of the *aapA3*/IsoA3 locus in an *E. coli* vector was essential to avoid contamination with WT *H. pylori* gDNA of the PCR products used in the transformation assays. To this end, the *aapA3*/IsoA3 locus was split into two fragments amplified with the Phusion High-Fidelity Hot Start DNA Polymerase and the primer pairs FA406/FA386 (“Up” fragment of 638 nt containing 415 nt of homology region, the *aapA3* promoter and the first 10 codons of the AapA3 ORF, Figure 1 – figure supplement 1) and FA409/FA387 (“Down” fragment of 680 nt containing IsoA3 promoter, the rest of aapA3 mRNA and 418 nt of homology region, Figure 1 – figure supplement 1). Note that the FA386 and FA387 primers have 25 nucleotides of overlap to allow PCR assembly. Each fragment was cloned in a separate pGEM®-T (Promega) plasmid (Table 2) and transformed into One-Shot TOP10 chemically competent *E. coli* cells (see Experimental Models in the KEY RESOURCES TABLE).

KEY RESOURCES TABLE.

KEY RESOURCES TABLE
| REAGENT or RESOURCE | SOURCE | IDENTIFIER |
| --- | --- | --- |
| **Chemicals and Recombinant Proteins** |  |  |
| DreamTaq DNA Polymerase | ThermoFischer Scientific | Cat#EP1701 |
| Phusion High-Fidelity DNA Polymerase | ThermoFischer Scientific | Cat#F530S |
| PfuUltra High-Fidelity DNA Polymerase | Agilent | Cat#600380 |
| Alkaline Phosphatase, Calf Intestinal (CIP) | New England Biolabs | Cat#M0290S |
| T4 Polynucleotide Kinase | New England Biolabs | Cat#M0201S |
| RNase T1 | New England Biolabs | Cat#AM2283 |
| *E. coli* RNase H1 | New England Biolabs | Cat#AM2293 |
| RNasin Ribonuclease Inhibitors | Promega | Cat#N2511 |
| **Recombinant DNA** |  |  |
| See Table 2 for the full list of plasmids used in this study | This paper | N/A |
| **Bacterial Strains I: *Escherichia coli*** |  |  |
| See Table 3 for the full list of *E. coli* strains used in this study | This paper | N/A |
| **Bacterial Strains II: *Helicobacter pylori*** |  |  |
| See Table 1 for the full list of *H. pylori* strains used in this study | This paper | N/A |
| **Critical Commercial Assays** |  |  |
| pGEM-T Easy Vector System | Promega | Cat#A1360 |
| MEGAScript T7 Kit | ThermoFischer Scientific | Cat#AM1334 |
| MAXIScript T7 Transcription Kit | ThermoFischer Scientific | Cat#AM1213 |
| High Purity Plasmid Miniprep Kit | Neo Biotech | Cat#NB-03-0002 |
| Quick Bacteria Genomic DNA extraction Kit | Neo Biotech | Cat#NB-03-0020 |
| *E. coli* 30S Extract System for Linear Templates Kit | Promega | Cat#L1030 |
| **Deposited Data** |  |  |
| Project and study description | This paper | NCBI BioProject PRJNA497299 |
| Deep-sequencing datasets raw data | This paper | NCBI SRA SRP166021 |
| Single-nucleotide substitutions, number of counts | This paper | NCBI GEO GSE121423 |
| Single-nucleotide substitutions, statistical analysis by sequence | This paper | NCBI GEO GSE121423 |
| Single-nucleotide substitutions, statistical analysis by position | This paper | NCBI GEO GSE121423 |
| Single-nucleotide deletions, statistical analysis by position | This paper | NCBI GEO GSE121423 |
| Single-nucleotide insertions, statistical analysis by position | This paper | NCBI GEO GSE121423 |
| **Experimental Models: Organisms/Strains** |  |  |
| *E. coli*: One Shot TOP10 chemically competent cells | ThermoFischer Scientific | Cat#C404010 |
| **Software and Algorithms** |  |  |
| Cutadapt 1.1 | DOI:10.14806/ej.17.1.200 | https://cutadapt.readthedocs.org/ |
| cmpfastq | NIHR Biomedical Research Centre for Mental Health | http://compbio.brc.iop.kcl.ac.uk/software/cmpfastq.php |
| Prinseq-lite 0.20.4 | (Schmieder and Edwards 2011) | http://prinseq.sourceforge.net/ |
| PANDaseq 2.9 | (Masella *et al.* 2012) | https://github.com/neufeld/pandaseq |
| BWA-SW algorithm of BWA 0.7.12 | (Li and Durbin 2009) | https://sourceforge.net/projects/bio-bwa/ |
| Samtools 1.2 | (Li *et al.* 2009) | https://sourceforge.net/projects/samtools/ |
| Bamtools 2.3.0 | (Barnett *et al.* 2011) | https://github.com/pezmaster31/bamtools |
| R 3.2.0 | (R Core Team 2015) | http://www.R-project.org/ |
| Differential analyses and Hierarchical tree clustering: Trinity 2.2.0 | (Haas *et al.* 2013) | http://trinityrnaseq.github.io |
| Differential analyses: DEseq2 1.10.1 | (Love *et al.* 2014) | http://www.bioconductor.org/packages/release/bioc/html/DESeq2.html |
| **Oligonucleotides** |  |  |
| See Table 4 for the full list of oligonucleotides used in this study | This paper | N/A |

### Mutant generation by Site-Directed mutagenesis PCR

Plasmids and custom-designed overlapping oligonucleotides containing the desired mutations were used for site-directed mutagenesis PCR using the PfuUltra high-fidelity DNA polymerase. To inactivate the IsoA3 −10 box, two synonymous point mutations (adenines +87 and +90 from the toxin TSS were mutated to cytosine and guanine, respectively) were introduced using the primer pair FA283/FA284 see Figure 1 for details). This strategy allowed us to preserve the toxin coding sequence while completely abolishing the transcription of the antitoxin, as previously shown (Arnion et al., 2017). To inactivate the toxin start codon, a single point mutation in the third codon position was introduced (thymine 54 was mutated to guanine) using the primer pair FA281/FA282 (see Figure 1 for details). WT or mutated fragments were amplified from the previously generated plasmids using the same primer pairs as those used for insert amplification prior to cloning. PCR assembly with 35 amplification cycles, the Phusion High-Fidelity Hot Start DNA Polymerase and the external primers FA406/FA409 was performed to generate the *aapA3*/IsoA3 locus variants (1294-nt amplicon) that were subsequently used as DNA substrates for *H. pylori* natural transformation. For the *in vivo* validation of the suppressor mutants studied here, the same protocol was used adapting the DNA oligonucleotides containing the desired mutations.

### Determining *H. pylori* transformation efficiency

For the transformation assays aiming at the determination of the transformation efficiency as an indirect proof of the toxicity of the expression of a PCR construct, transformation patches (after 16 hours growth upon DNA addition) were recovered and resuspended in 1 mL BHI. Ten-fold serial dilutions adapted to each transformation case (10^7^, 10^6^ and 10^5^ for non-selective media; and 10^4^, 10^3^, 10^2^ for selective media upon transformation with water or a toxic construct; and 10^5^, 10^4^, 10^3^ for selective media upon transformation with non-toxic constructs) were performed. Allelic replacement events were selected by the use of streptomycin-containing plates (selection of loss of the *rpsl_Cj_-erm* cassette, Str^R^). The number of Str^R^ CFU/ total CFU was calculated, plotted and statistically analyzed by unpaired *t* (student)-test (GraphPad Prism software version 7).

### *H. pylori* transformation assay to identify toxicity suppressors by Illumina sequencing

Transformation assays to select toxicity suppressors were performed in three biological replicates using the wild-type (WT) or antitoxin promoter inactivated PCR-generated constructs (pIsoA3*). Upon transformation, all bacteria were recovered and serially diluted. Transformants were selected on streptomycin-containing plates by using optimized dilutions (9 plates/replicate of 10^1^ dilution for pIsoA3* and 3 plates/replicate of 10^3^ dilution for WT). Three days after transformation, colonies were pooled (approximately 60,000 colonies per transformation) and genomic DNA was extracted. Next, the *aapA3*/IsoA3 locus was amplified with the primer pair FA395/FA396 (426-nt amplicon, Figure 1 – figure supplement 1), which allows the introduction of the DNA adapters for Illumina paired-end sequencing. Importantly, to avoid amplification from phenotypic revertant clones (mutated in the *rpsL* gene), the FA395 and FA396 primers are nested to the ones used for locus deletion (FA407 and FA408, Figure 1 – figure supplement 1), thus, binding to deleted regions that are re-introduced only upon recombination. For this PCR, the Phusion High-Fidelity Hot Start DNA polymerase (Thermo Fisher) and 35 amplification cycles were used. Finally, the samples were sent for sequencing at the Platforme GeT-PlaGe-, Genotoul Centre INRA, Toulouse, France. Sequencing was done on an Illumina MiSeq instrument in paired-end mode 2 × 250 nt (overlapping reads).

### Polysome fractionation in sucrose gradients

*H. pylori* strains were grown as described above. At an early exponential phase (OD_600nm_<0.9), chloramphenicol (100 µg.mL-1) was added to the culture to stabilize translating ribosomes. After 5 min incubation at 37°C, cultures were quickly cooled by transferring them into pre-chilled flasks immerged in a dry ice/ethanol bath. Cultures were then centrifuged for 10 min at 3,500 rpm and 4°C and pellets were washed with Buffer A (10 mM Tris-HCl pH 7.5; 60 mM KCl; 10 mM MgCl_2_) and frozen at −80°C. Then, pellets were resuspended in 500 µl of Buffer A containing RNasin® Ribonuclease Inhibitor (Promega) and cells were lyzed with glass beads in a Precellys homogenizer (Bertin). Lysates were recovered and immediately frozen in liquid nitrogen. About 10 OD_260_ units of lysate were layered onto 10%-40% sucrose gradients in Grad-Buffer (10 mM Tris-HCl pH 7.5; 50 mM NH_4_Cl; 10 mM MgCl_2_; 1mM DTT) and centrifuged at 35,000 rpm for 3.75 h at 4°C in a SW41 Ti rotor. Gradients were analysed with an ISCO UA-6 detector with continuous OD monitoring at 254 nm. Fractions of 500 µl were collected and RNA was precipitated overnight at −20°C in the presence of 1 volume of ethanol containing 150 mM of sodium acetate (pH 5.2). RNA was extracted and subjected to Northern Blot analysis following the protocols described above.

## BIOINFORMATIC AND STATISTICAL NGS DATA ANALYSES

### Read pre-processing and alignment

Reads were first trimmed of low-quality 3’ ends using cutadapt 1.1 (https://cutadapt.readthedocs.org/) and a base quality threshold of 28 (option “-q 28”). Then, reads having an average base quality lower than 28 were discarded using prinseq-lite 0.20.4 (“-min_qual_mean 28”; (Schmieder & Edwards, 2011)). Read pairs for which both mates passed the quality filtering steps were recovered by means of cmpfastq (http://compbio.brc.iop.kcl.ac.uk/software/cmpfastq.php), and mates were assembled into a single sequence using PANDAseq 2.9 (Masella, Bartram, Truszkowski, Brown, & Neufeld, 2012) run with options “-N -o 30 -O 0 -t 0.6 -A simple_bayesian -C empty”. About 5 million read pairs (combining the three biological replicates) could be assembled for the WT (*aapA3*/pIsoA3) and pIsoA3* (*aapA3*/pIsoA3*) samples. These assembled reads were aligned onto the 426-nt reference sequence by the BWA-SW algorithm of BWA 0.7.12 (Li & Durbin, 2009) run with options “-a 1 -b 3 -q 5 -r 2 -z 1” to produce alignments in BAM format. Mapped sequences of length 426 showing a single substitution compared to the reference were then extracted using utilities from the samtools 1.2 (Li et al., 2009) and bamtools 2.3.0 (Barnett, Garrison, Quinlan, Stromberg, & Marth, 2011) packages based on the various flags and tags in the BAM files (in particular the CIGAR string and NM tag). This gave a dataset of 1,653,406 WT and 2,559,164 pIsoA3* single-substitution sequences. Mapped sequences of length 425 and 427 harboring a single deletion or insertion, respectively, were also extracted (40,998 WT and 100,754 pIsoA3* single-deletion sequences; 4,799 WT and 7,048 pIsoA3* single-insertion sequences).

### Statistical analysis

Statistical analyses of the differential distribution of substitutions in the WT and pIsoA3* single-substitution sequences were carried out. To determine whether substitutions were enriched at particular positions in the pIsoA3* compared to WT sequences, a “positional” analysis was conducted by summing together the counts of all sequences that showed a substitution at a given position, regardless of the identity of the substituted nucleotide. A “nucleotide-specific” analysis comparing the amount of each individual sequence was also done to determine whether particular nucleotides were enriched at specific positions. As positions +87 and +90 were mutated to inactivate the IsoA3 promoter (see Figure 1B), for the “nucleotide-specific” analysis all sequences showing a difference to the reference at one or both of these two positions were excluded from the pIsoA3* and WT datasets (11,319 pIsoA3* and 38,575 WT sequences, respectively), and the pIsoA3* reference was converted back to the WT reference in order to make data comparable between WT and pIsoA3* samples. Differential analyses were conducted following the protocol of Haas *et al*. (Haas et al., 2013) using tools from the Trinity 2.2.0 (Haas et al., 2013) and DESeq2 1.10.1 packages (Love, Huber, & Anders, 2014), taking into account variability among the three biological replicates. The sequence abundance estimation step was not performed; actual sequence counts were used. The four positions at each extremity of the 426-nt amplicon corresponding to pieces of the primers could not be included in the statistical analyses as there were no substitutions at these positions in any the WT and pIsoA3* replicates. Substitutions were considered as significantly over- or under-represented in the pIsoA3* vs. WT samples if the *p*-value adjusted for multiple testing (False Discovery Rate [FDR] calculated using the Benjamini-Hochberg [BH] method in DESeq2) was equal or lower than 5% (padj ≤ 0.05). Bar plots of normalized sequence counts and log2 ratios of fold change were drawn using R 3.2.0 (R Core Team, 2015. R: A language and environment for statistical computing. R Foundation for Statistical Computing, Vienna, Austria; http://www.R-project.org/). Similar “positional” analyses were also carried out for single-insertion and single-deletion sequences to determine whether insertions or deletions were statistically enriched at particular positions in the pIsoA3* dataset.

For the “positional” analysis, a heatmap and hierarchical tree clustering of samples according to sequence count patterns was also performed. This was based on TMM-normalized (trimmed mean of M values), median-centered, log2-transformed FPKM (fragment per kilobase per million reads mapped) values, computed according to the protocol and tools of Haas *et al*. (Haas et al., 2013). The Pearson correlation coefficient was used as distance metric and average linkage was chosen as clustering method (options “--sample_dist sample_cor --sample_cor pearson --sample_clust average” for the “analyze_diff_expr.pl” utility script). A log2 cut-off of 0 and a p-value cut-off of 1 were set (options “-C 0 -P 1”) in order to include all sequence positions in the map. The clustering script “PtR” was manually edited to suppress the clustering by sequence (*i.e.*, rows) and sort the positions by numerical order instead.

## DATA AVAILABILITY

The deep-sequencing raw and analyzed datasets reported in this paper have been deposited in the National Center for Biotechnology Information Gene Expression Omnibus (NCBI GEO) data repository under the accession code GSE121423 and can be accessible at the URL: https://www.ncbi.nlm.nih.gov/geo/query/acc.cgi?acc=GSE121423

## ACKNOWLEDGMENTS

We thank all present and past members of the ARNA laboratory for helpful discussions and in particular, Anais Le Rhun for critical reading of the manuscript. This work was supported by INSERM U1212, CNRS UMR 5320, Université de Bordeaux, and Agence Nationale de la Recherche (http://www.agence-nationale-recherche.fr/) grants Bactox1 and asSUPYCO. This work was performed in collaboration with the GeT core facility, Toulouse, France (http://get.genotoul.fr), and was then supported by France Génomique National infrastructure, funded as part of “Investissement d’avenir” program managed by Agence Nationale pour la Recherche (contract ANR-10-INBS-09). This project has also received funding from the European Union’s Horizon 2020 research and innovation programme under the Marie Sklodowska-Curie grant agreement No 642738. The funders had no role in study design, data collection and analysis, decision to publish, or preparation of the manuscript.

---

**Figure 1–figure supplement 1.**
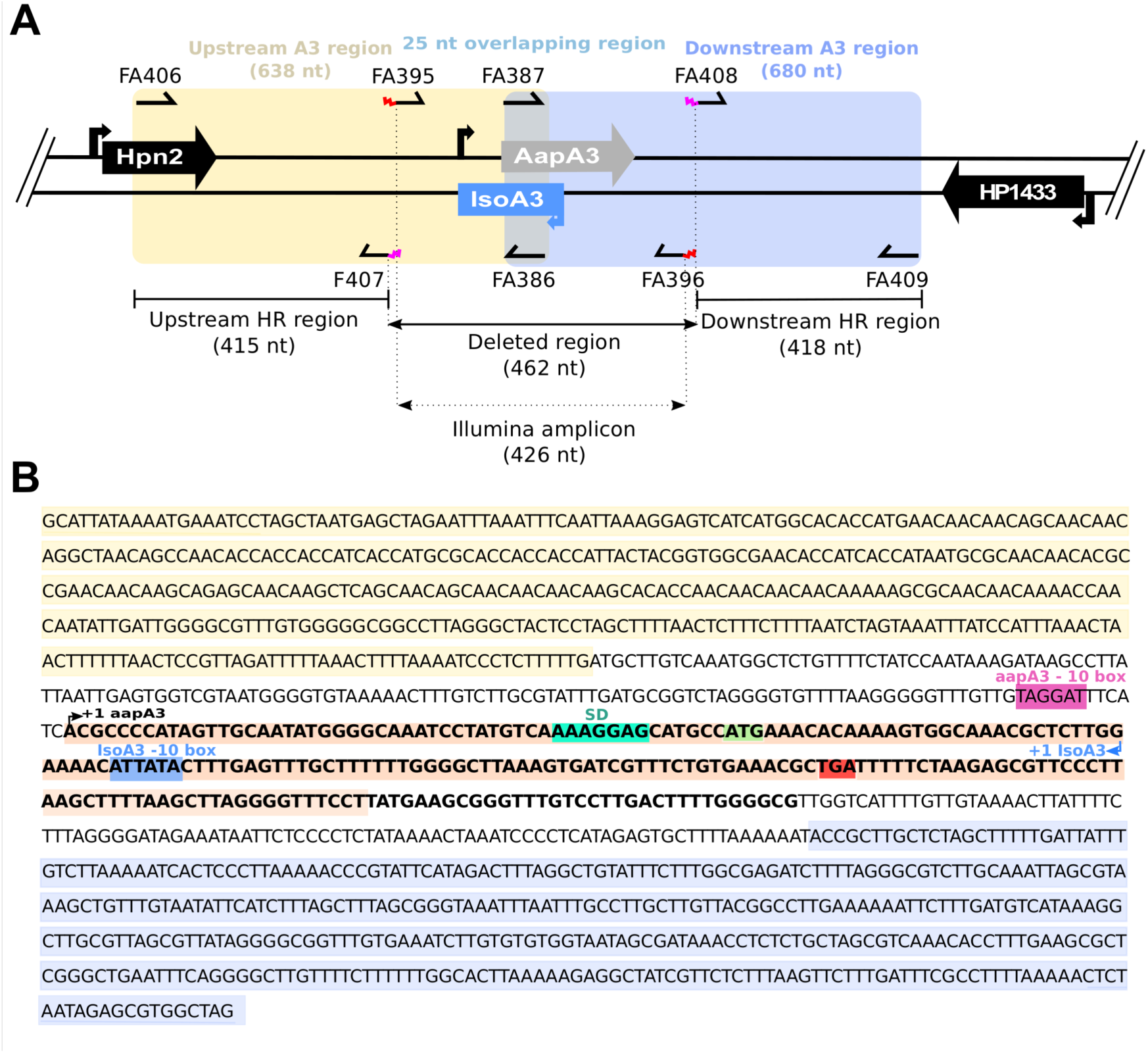
Details on *aapA3*/IsoA3 locus deletion and deep-sequencing approaches. **(A)** Schematic representation of the oligonucleotides used for the *aapA3*/IsoA3 locus deletion and Illumina paired-end sequencing. Are shown: the upstream and downstream homology regions (HR) used for homologous recombination in all transformation assays; the deleted region and the amplicon used for Illumina paired-end sequencing approach; primers FA407 and FA408 carrying an overhang 5’ tail (in pink) containing the 5’ and 3’ ends, respectively, of the *rpsl_Cj_-erm* cassette used for locus deletion; primers FA395 and FA396 carrying an overhang 5’ tail (in red) containing the adapters for paired-end Illumina sequencing approach (see Table S4 for oligonucleotides sequences). **(B)** The *aapA3*/IsoA3 module located at the chromosomal locus III of the *H. pylori* 26695 strain. Upstream (415 nt) and downstream (418 nt) regions used for homologous recombination (HR) are highlighted in yellow and purple, respectively. Deleted region in the *aapA3*/IsoA3 knockout mutant corresponds to the region flanked by the upstream and downstream HR regions. AapA3 −10 box is shown in pink. AapA3 transcription start site (TSS, +1 *aapA3*) determined by RNA-seq analysis is represented with a black arrow. AapA3 full-length transcript is highlighted in bold. 3’ end-truncated *aapA3* mRNA is highlighted in orange. AapA3 Shine-Dalgarno sequence (SD) is shown in turquoise. AapA3 start (ATG) and stop (TGA) codons are shown in green and red, respectively. IsoA3 −10 box is highlighted in blue. IsoA3 transcription start site (TSS, +1 IsoA3) is represented by a blue arrow.

**Figure 3–figure supplement 1.**
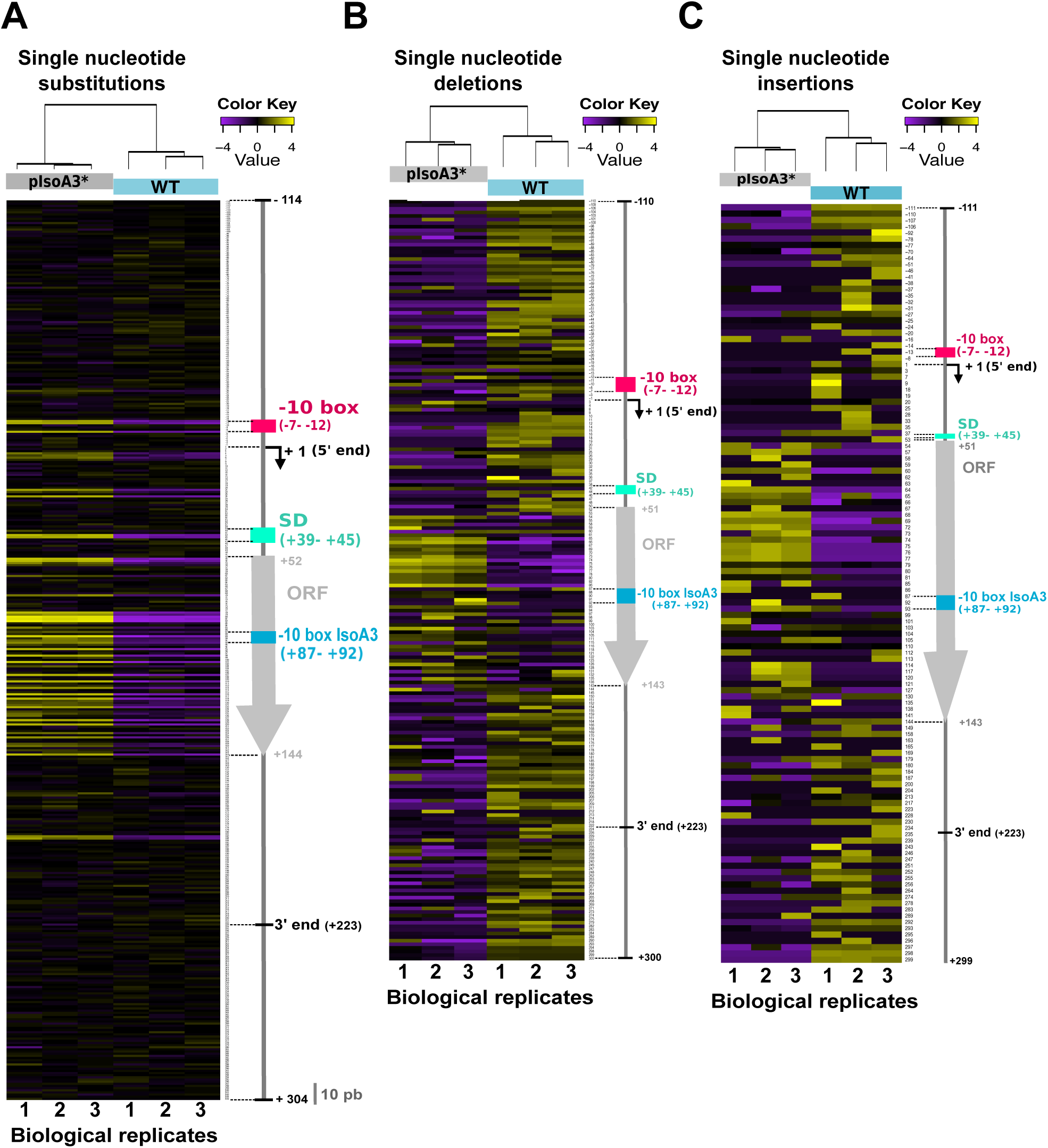
Comparison of the distribution and relative frequency of single-nucleotide suppressors in the WT and pIsoA3* *aapA3*/IsoA3 modules. Heatmap of the location and relative abundance of single-nucleotide substitutions **(A)**, deletions **(B)**, and insertions **(C)**, in WT (*aapA3*/IsoA3) and pIsoA3* (*aapA3*/pIsoA3*) samples coming from Miseq Ilumina deep-sequencing. For a given position, all sequences showing a single-nucleotide substitution **(A)**, deletion **(B)**, or insertion **(C)** at that position were counted irrespective of the nucleotide identity (“positional” analysis). Sequence counts were converted into TMM-normalized, median-centered, and log2-transformed FPKM values, which were then hierarchically clustered according to biological replicate sample (see Methods section for details). The color key gives the log2 value scale (negative and positive values represent relative frequencies below and above the median, respectively). Positions are numbered relative to the *aapA3* transcriptional start site (TSS, +1). Colored arrows and boxes on the right-hand side of each heatmap indicate the locations of relevant sequence elements on the *aapA3/*IsoA3 locus: −10 box, *aapA3* promoter −10 box; −10 box IsoA3, IsoA3 promoter −10 box; +1 (5’ end), *aapA3* transcription start site; SD, Shine-Dalgarno; ORF, open reading frame; 5’ and 3’ ends delimit the UTRs (untranslated regions) on the *aapA3* mRNA; the “10 nt” scale bar at the bottom of panel (A) is used to measure intervals of 10 nucleotides alongside the map.

**Figure 3–figure supplement 2.**
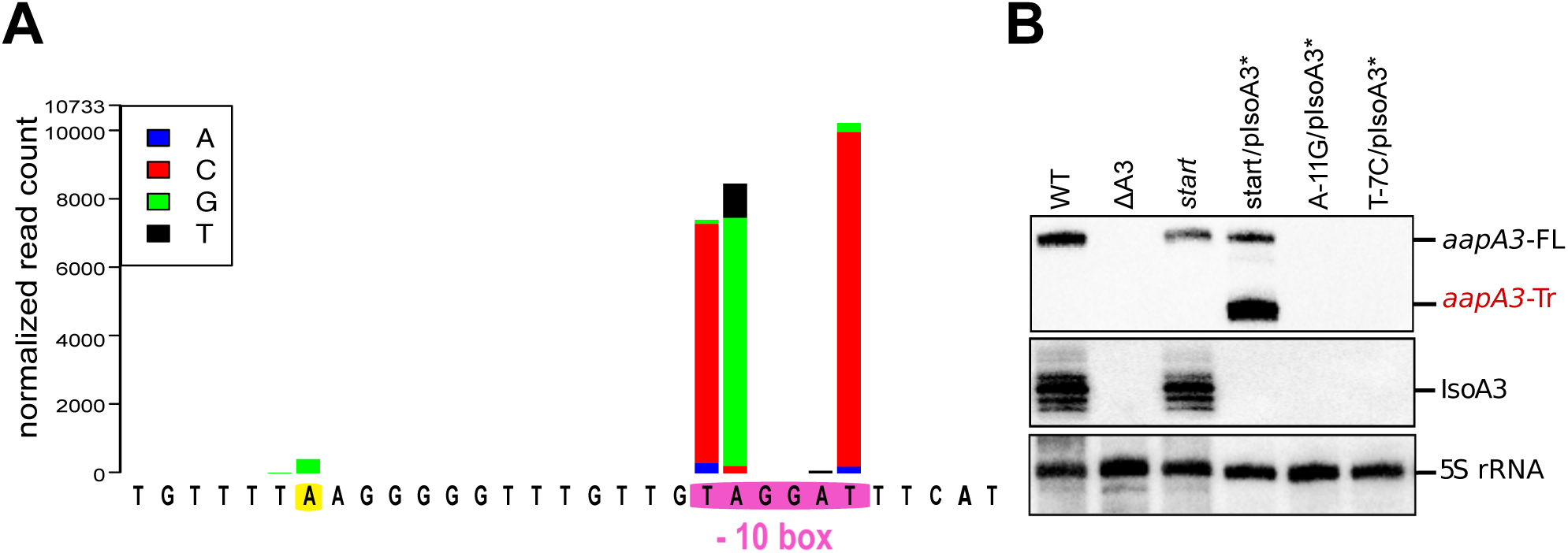
Defining and validating AapA3 promoter with nucleotide resolution. **A)** Statistical analysis of the differential amount of individual single-nucleotide substitution sequences in WT (*aapA3*/IsoA3) and pIsoA3* (*aapA3*/pIsoA3*) samples was carried out using DESeq2 (“nucleotide-specific” analysis; see Methods section for details). Nucleotide substitutions that are significantly enriched (padj≤0.05) in pIsoA3* compared to WT sequences in *aapA3* promoter region are shown. The −10 box of the *aapA3* gene promoter is highlighted in pink; an enriched mutation located at position −26 from *aapA3* +1 is highlighted in yellow. **(B)** Total RNA was isolated from the indicated strains and 10 µg were subjected to Northern Blot analysis. The same membrane was successively probed with FD38 labeled oligonucleotide and IsoA3 riboprobe to detect *aapA3* and IsoA3 transcripts, respectively. The different transcripts are annotated as: *aapA3*-FL (full length), *aapA3*-Tr (3’ end truncated), and IsoA3 full-length and processed transcripts. Proper loading was assessed by the level of 5S rRNA using the labeled oligoprobe FD35.

**Figure 4–figure supplement 1.**
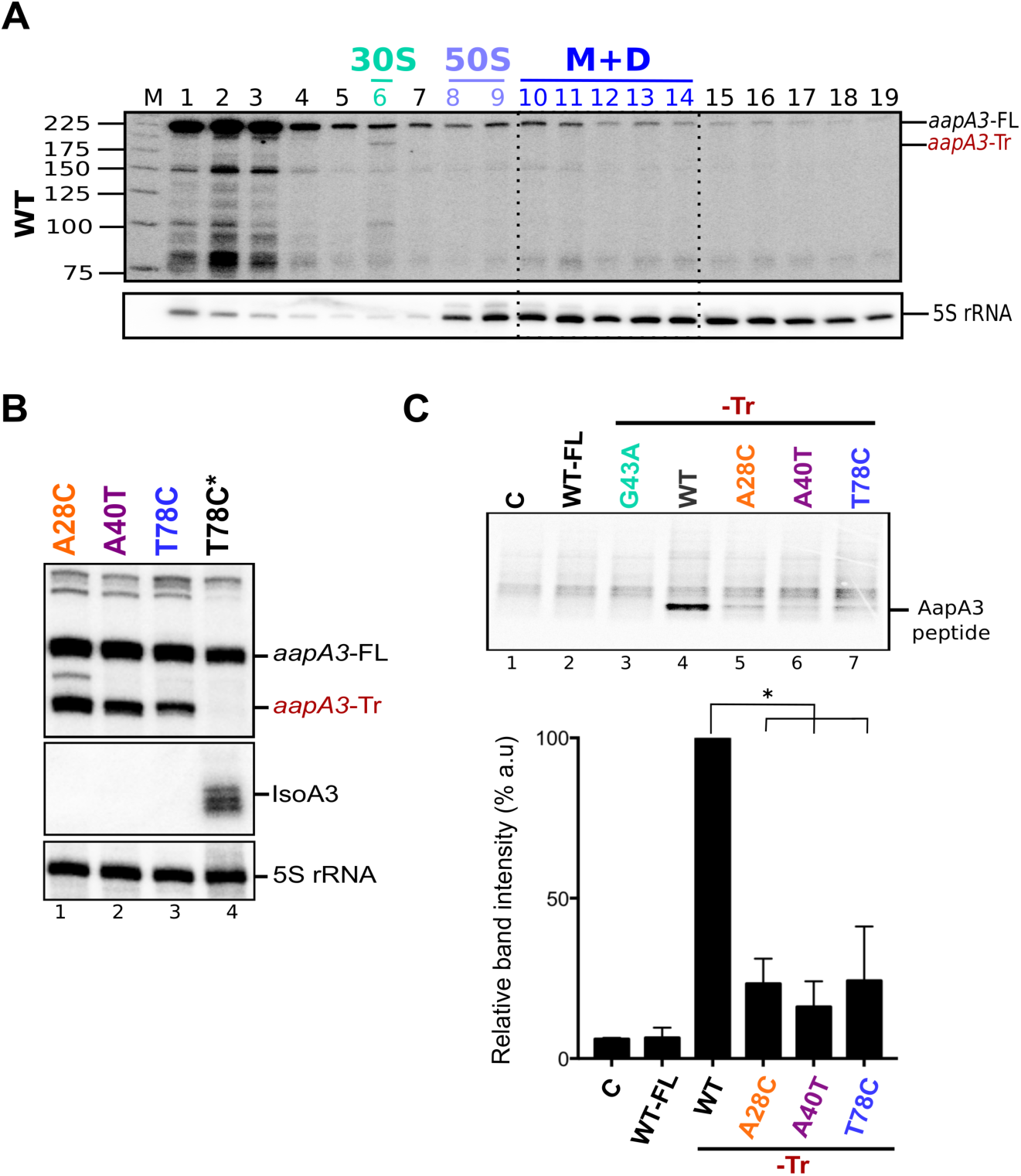
The A28C, A40T and T78C suppressors inhibit *aapA3*-Tr mRNA translation. **(A)** Only the 3’ end truncated *aapA3* mRNA form is translated *in vivo*. Cell lysate of the *H. pylori* 26695 wild-type (WT) strain was subjected to ultracentrifugation through a sucrose gradient under polysome stabilization conditions (+ Chloramphenicol). A profile at OD_254nm_ was recorded. RNA was extracted from each fraction and equal volumes of each extract were subjected to Northern Blot analysis. Fractions corresponding to the free 30S and 50S subunits, 70S ribosomes (free and translating) and polysomes are indicated. The different transcripts are annotated as: *aapA3*-FL (full length, 225 nt), *aapA3*-Tr (3’ end truncated, 190 nt), and 5S rRNA as loading control (5S rRNA). M, monosomes; D, disomes. **(B)** Gene expression analysis of the indicated strains was analyzed by Northern Blot. Transcripts *aapA3*-FL (full length), *aapA3*-Tr (3’ end truncated), and IsoA3 full-length and processed transcripts are shown. Proper loading was assessed by the level of 5S rRNA. The T78C* construct contains a WT IsoA3. **(C)** Relative peptide production upon *in vitro* translation of *aapA3*-FL, *aapA3*-Tr and the *aapA3*-Tr form of the three independent suppressor mutants A28C-Tr, A40T-Tr and T78C-Tr (upper panel). A construct with inactivated SD sequence (G43A-Tr) was also included for comparison. Control lane (0) shows the translation background obtained without exogenous mRNA. A representative experiment is shown. Relative peptide production was quantified (lower panel). Error bars represent the s.d; *n=3* technical replicates, \**P*<0.0001 according to unpaired t-test. Peptide production using the G43A-Tr control RNA was not quantified as the experiment was performed only once. Figure 4–figure supplement 1-source data 1

**Figure 4–figure supplement 2.**
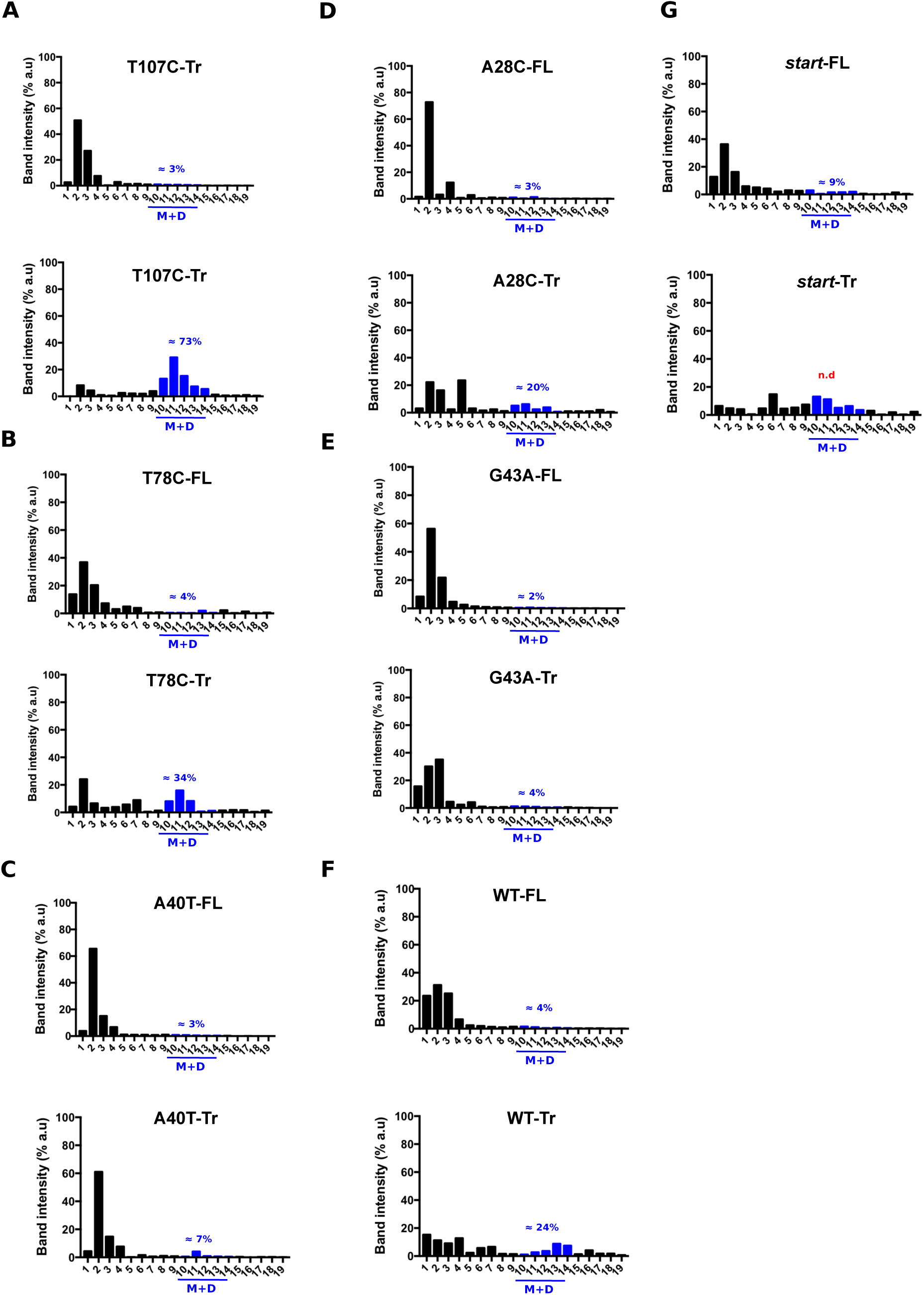
Quantification of the relative *aapA3* mRNA band intensity from polysome fractionation Northern Blots shown in Figure 4. Relative band intensity was determined using the ImageLab software. Percentage of band intensity located on 70S fractions was calculated for each *aapA3* transcript (full-length, -FL or truncated, -Tr) by dividing the band intensity in 70S fractions by the total band intensity (intensity values in all fractions). Each panel shows the data for a given WT or mutant transcript. M, monosomes; D, disomes. In the case of the *start* (G54T/pIsoA3*) strain, the quantification of the *aapA3*-Tr form present on the M+D fractions was not possible due to a strong mRNA degradation (n.d= non-detectable).

**Figure 5–figure supplement 1.**
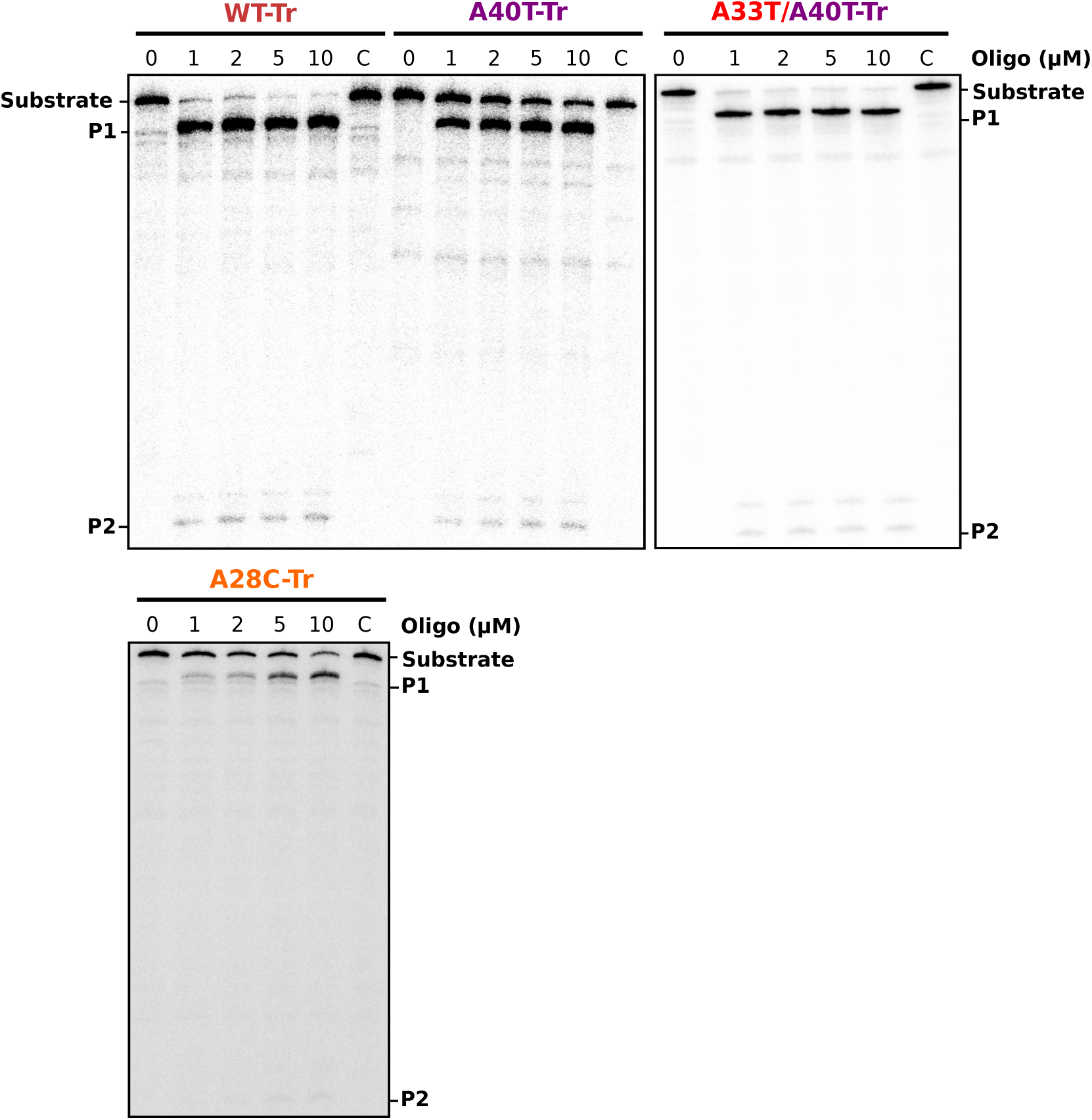
Gel analysis of RNase H/ oligonucleotide accessibility assays. 30 fmol of internally labeled RNA were used. DNA oligonucleotides were used to a final concentration of 0 to 10 µM. Reactions were incubated for 30 min at 30°C in the presence or absence (C, control) of 0.25 U *E. coli* RNase H1. Reactions were stopped with 10 µl of 2X Loading Buffer and products were analyzed in an 8% denaturing PAA gel. See Figure 5B for relative band quantification. P1 and P2 indicate the two RNase H-oligonucleotide-mediated RNA cleavage products.

**Figure 6–figure supplement 1.**
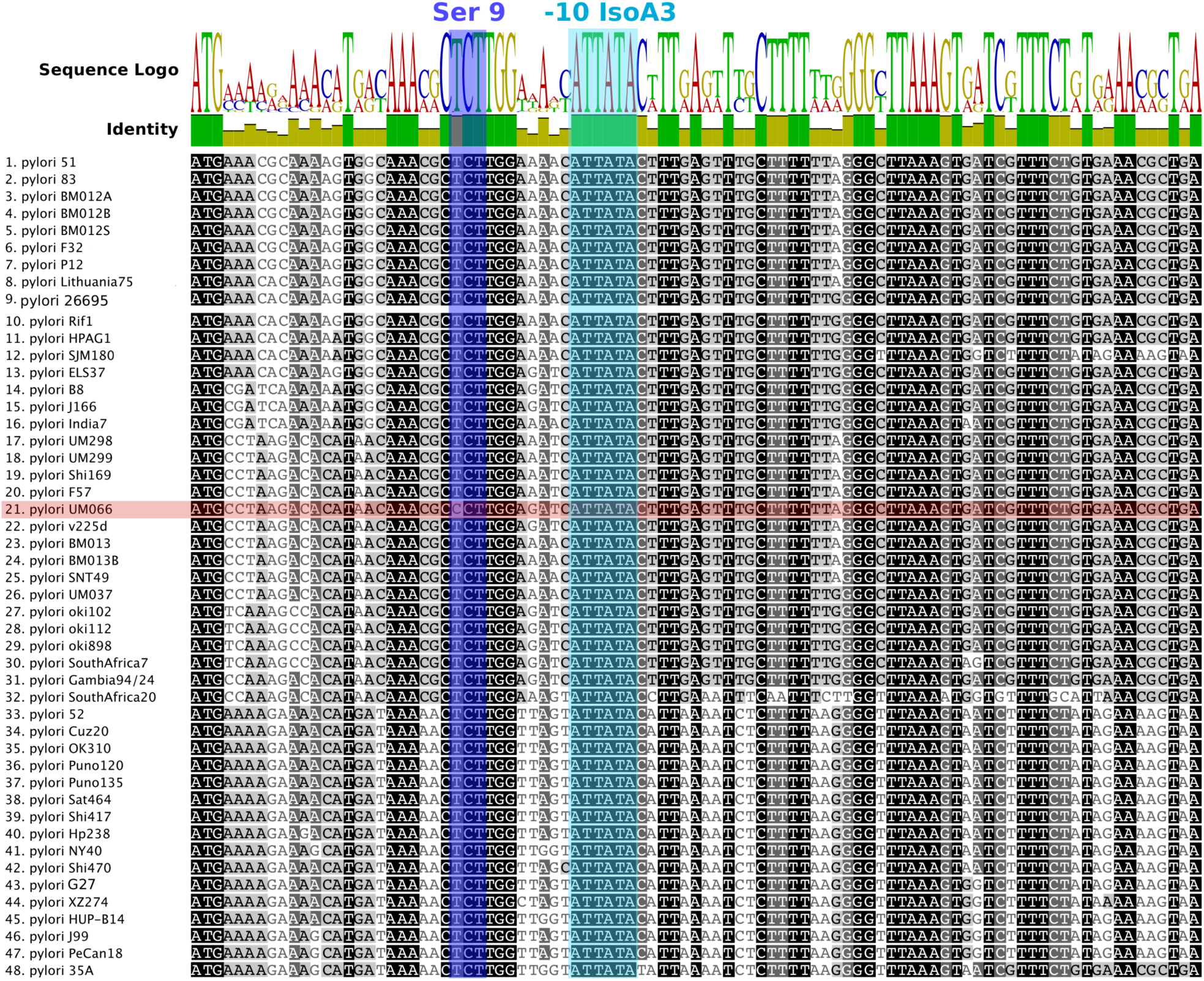
Nucleotide alignment of AapA3 coding region of 49 *Helicobacter pylori* strains. Conserved nucleotides are highlighted in different tones of grey depending on their conservation level. Sequence logo and identity scores are shown on the top. The highly conserved region corresponding to the second SD sequestering-sequence (serine residue at position 9) and the IsoA3 −10 box are highlighted in dark and light blue, respectively. The UM066 strain, only strain containing a CCT Proline codon at position 9, is highlighted in pink. Geneious software 8.1.8 (Kearse et al., 2012) was used for sequence collection and alignment.

**Figure 7–figure supplement 1.**
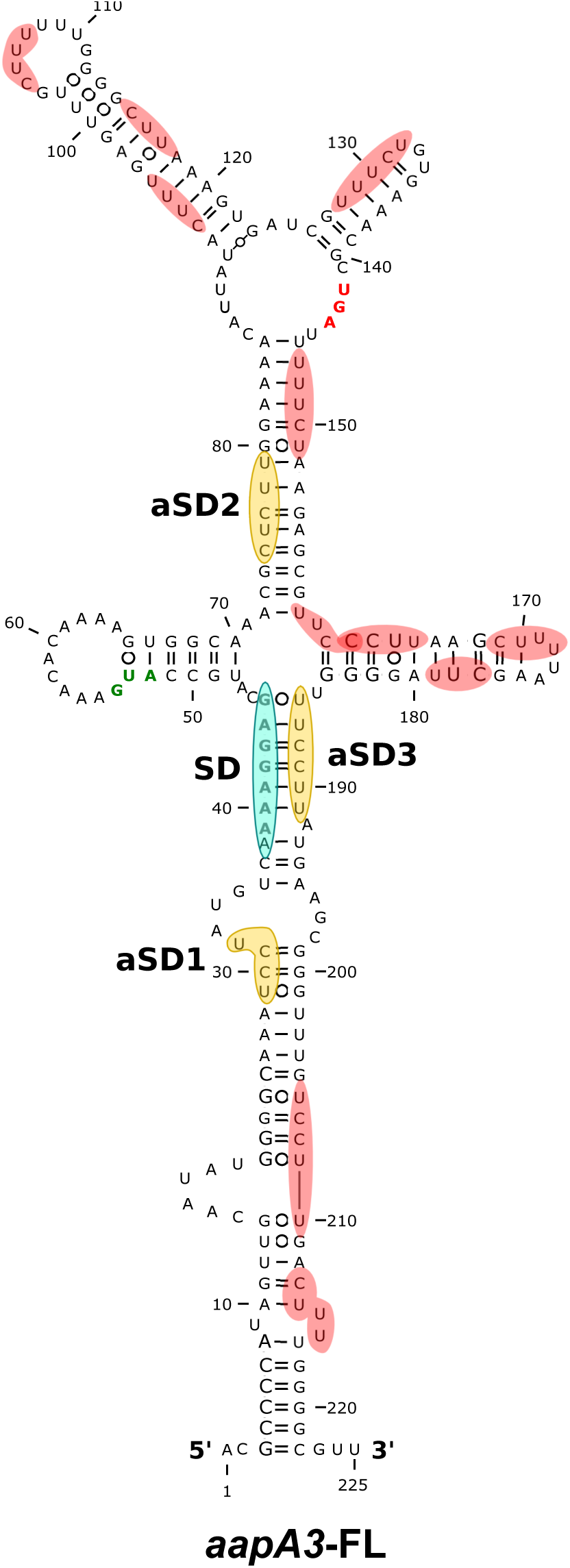
Only three out of the thirteen potential aSD sequenced embedded in *aapA3* mRNA are functional. 2D structure predictions were generated with the RNAfold Web Server (Gruber et al., 2008) and VARNA (Darty et al., 2009) was used to perform the drawing. Potential, but not used, anti-Shine-Dalgarno (aSD) sequences (UC-rich motifs) are highlighted in red. The three functional aSD sequences (aSD1, aSD2, aSD3) are shown in gold. Shine-Dalgarno (SD) sequence is shown in turquoise. Translation start and stop codons are shown in green and red, respectively.

**Figure 7–figure supplement 2.**
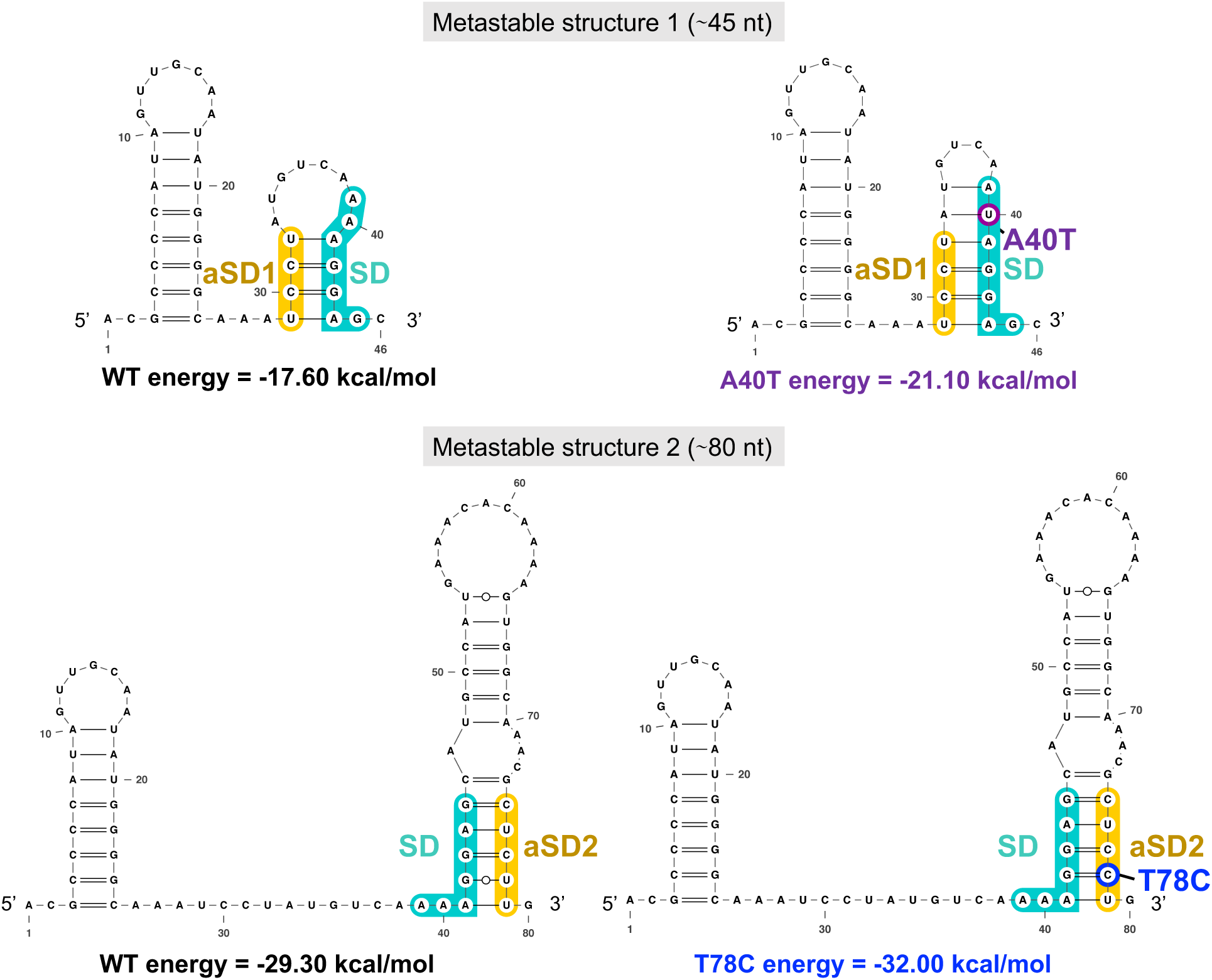
The two successive AapA3 mRNA metastable structures have increasing stability and are stabilized by the A40T and the T78C suppressors. The two metastable structures (1, ≈46-nt long, upper panel; and 2, ≈80-nt long, lower panel) successively formed during AapA3 mRNA transcription are shown. Shine-Dalgarno (SD) sequence is highlighted in turquoise, anti-SD (aSD) sequences (aSD1 and aSD2) are highlighted in yellow, suppressor mutation stabilizing SD sequestration by aSD1 in the metastable structure 1 is highlighted in purple (A40T), the suppressor stabilizing the SD sequestration by aSD2 in the metastable structure 2 is highlighted in blue (T78C). RNAfold (Gruber et al., 2008) was used for secondary structure and minimum free energy predictions, and VARNA (Darty et al., 2009) for drawing.

